# Hepatitis C virus non-structural proteins modulate cellular kinases for increased cytoplasmic abundance of host factor HuR and facilitate viral replication

**DOI:** 10.1101/2022.08.08.503121

**Authors:** Harsha Raheja, Biju George, Sachin Kumar Tripathi, Sandhini Saha, Tushar Kanti Maiti, Saumitra Das

## Abstract

Host protein HuR translocation from nucleus to cytoplasm following infection is crucial for the life cycle of several RNA viruses including hepatitis C virus (HCV), a major causative agent of hepatocellular carcinoma. HuR assists the assembly of replication-complex on the viral-3′UTR, and its depletion hampers viral replication. Although cytoplasmic HuR is crucial for HCV replication, little is known about how the virus orchestrates the mobilization of HuR into the cytoplasm from the nucleus. We show that two viral proteins, NS3 and NS5A, act co-ordinately to alter the equilibrium of the nucleo-cytoplasmic movement of HuR. NS3 activates protein kinase C (PKC)-δ, which in-turn phosphorylates HuR on S318 residue, triggering its export to the cytoplasm. NS5A inactivates AMP-activated kinase (AMPK) resulting in diminished nuclear import of HuR through blockade of AMPK-mediated phosphorylation and acetylation of importin-α1. Cytoplasmic retention or entry of HuR can be reversed by an AMPK activator or a PKC-δ inhibitor. Our findings suggest that efforts should be made to develop inhibitors of PKC-δ and AMPK, either separately or in combination, to inhibit HCV infection.

**Author summary:** Hepatitis C virus is a major human pathogen, which exploits cellular machinery for its propagation in liver cells. The cytoplasmic availability of cellular components is crucial for their direct influence on processes involving the viral RNA, which lacks any nuclear history. Our results establish the involvement of viral proteins, NS3 and NS5A in achieving increased cytoplasmic abundance of a host factor HuR, an RNA binding protein (RBP) critical for HCV replication. This is achieved via direct post-translational modification of HuR and indirect regulation of its nuclear carrier by coercing two host kinases, PKC-δ and AMPK-α. RBPs are emerging as novel targetable candidates for gene regulation. Similar studies with other RBPs and targeting protein modifications, in place of whole protein knockdown, could usher in a revolutionary strategy to neutralize emerging RNA virus-based diseases, while preserving their cellular functions.

## Introduction

Hepatitis C virus (HCV) is an RNA virus, whose 9.6-kb genome codes for three structural and seven non-structural proteins (1). Structural proteins form the viral capsid and envelope, whereas non-structural proteins are involved in the regulation of various viral life processes, including double-membrane vesicle formation, replication, translation, packaging, and release (2–5). The viral open reading frame is flanked by two regulatory elements, the 5’-untranslated region (UTR) and 3’-UTR, which are involved in initiating viral translation and replication, respectively (5).

Multiple host factors are required at various stages of the viral life cycle. Upon viral infection, proteins, such as some IRES trans-acting factors and Nups relocate from the nucleus to the cytoplasm to regulate the viral life cycle (6–8). Our laboratory has previously shown that human antigen R (HuR), also known as ELAVL1, relocates from nucleus to the cytoplasm upon HCV infection. HuR binds to the viral 3’-UTR, where it displaces the replication inhibitor protein PTB and assists in the binding of the La protein. La further recruits the entire replication complex to initiate viral replication (9). HuR is an RNA-binding protein (RBP), that shuttles between nucleus and cytoplasm. Its endogenous function is to regulate the transport, stability, and translation of cellular RNAs (10–12). Infection-induced relocalization of HuR to the cytoplasm is required for interaction with viral RNA, which guides viral replication. In a genome wide CRISPR screen, HuR was identified as a major host factor required for HCV replication and the specific knockout was found to abolish the HCV RNA replication in cells (13).

HuR contains three RNA recognition motifs (RRMs) and a hinge region, also known as the HuR nuclear shuttling domain (HNS domain) (14–16). Phosphorylation of HuR residues regulates its structure, RNA-binding affinity, and subcellular localization (17). HuR is phosphorylated in a cell-cycle dependent manner by CDKs. CDK1 phosphorylates HuR on S202, which is required for its nuclear retention during the S and G2 phases of the cell cycle (18). Phosphorylation on the same site by CDK5 has been shown to reduce its binding to cyclin A mRNA, leading to cell cycle arrest in glioma cells (19). Checkpoint kinase 2 (CHK2) phosphorylates HuR on S88, S100, and T118, which modulates HuR binding to RNA by decreasing its affinity for certain mRNAs (e.g., SIRT1) and increasing it for others (e.g., Myc and occludin) (20). Phosphorylation on S242 augments the amount of HuR in the cytoplasm, regardless of stress conditions such as exposure to short-wavelength ultraviolet light (21). The MAP kinase p38, which is activated during inflammatory response, phosphorylates HuR on T118, increasing its binding to and the stabilization of inflammatory mRNAs, such as p21, COX2, and cPLA_2_α, in addition to promoting HuR accumulation in the cytoplasm (22, 23). Several members of the protein kinase C (PKC) family are also involved in HuR phosphorylation. PKC-α phosphorylates HuR on S158 and S221, which leads to its cytoplasmic relocalization and enhanced binding to targets, such as COX2 and PTGS2 (24). PKC-δ-mediated phosphorylation on S318 and S221 has been shown to induce cytoplasmic translocation and increased binding to COX2, CCNDA2, and CCND1 mRNAs (25, 26). Abrogation of phosphorylation on S318 hinders the migration and invasion of colon cancer cells (27). Other kinases, such as IKKα, ERK8, and JAK3, also phosphorylate HuR to regulate its localization and function (28–30). Activated JAK3 phosphorylates HuR on Y200, which prevents its inclusion in stress granules formed upon arsenic treatment (28). Phosphorylation of Y5, Y95, Y105, and Y200 residues by SRC and ABL1 kinases is important for centrosomal accumulation of HuR and, therefore, genomic stability (31). In addition to phosphorylation, HuR is methylated on R217 by CARM1, which correlates with its cytoplasmic localization in non-small cell lung carcinoma (32). Ubiquitination on K313 and K326 leads to the dissociation of HuR from p21, MKP1, and SIRT1 mRNA (33); whereas neddylation on K283, K313, and K326 promotes nuclear localization and protein stability (34).

Subcellular localization of HuR is also regulated by AMP-activated kinase (AMPK) (35). Inhibition of AMPK has been associated with cytoplasmic localization of HuR, leading to increased stability of the HuR-bound transcripts p21, cyclin B1, cyclin A, and IL-20. AMPK phosphorylates importin-α1 on S105 and acetylates it on K22, thereby preventing its interaction with HuR and import to the nucleus (36).

Cells adopt multiple strategies to regulate the subcellular localization of HuR and, consequently, the stability of its cognate transcripts under different conditions. Here, we investigated the viral mechanism driving the nuclear export of HuR, the cytoplasmic partitioning of HuR, and the effect of its altered localization. We report that PKC-δ and AMPK work in a coordinated manner to achieve cytoplasmic localization of HuR via direct modification and indirect regulation of its nuclear carrier. Thus, HCV modulates two separate kinases to promote replication and pathogenesis. Relocalization of HuR and its interaction with cytoplasmic RNAs could reveal the molecular basis of HCV-induced hepatocellular carcinoma and provide an attractive target for preventing its progression.

## Results

### Phosphorylation of HuR regulates its cellular localization upon HCV infection

It has been shown that HuR relocalises from nucleus to cytoplasm after 48h of HCV infection and is important for HCV replication (9, 13). These studies have been done in Huh7.5 and Huh7.5.1 cells. Therefore, we first studied the time kinetics of this relocalisation using nuclear-cytoplasmic fraction of mock and HCV-JFH1 virus infected Huh7.5 cells at 12h and 24h post infection (Fig S1A, B). We calculated the ratio of nuclear to cytoplasmic HuR intensities and found that the increase in cytoplasmic HuR begins from 12h post infection and increases till 24h. These time points correlate with the replication phase of the virus wherein translation to replication switch occurs 12-18h post infection and highlight the importance of HuR relocalisation for HCV replication. Viruses are known to modify host proteins (37). To study the post-translational modifications of host proteins after HCV infection, total protein extracts prepared from mock (control) and HCV-RNA transfected Huh7.5 cells were analyzed by two-dimensional gel electrophoresis and HuR-specific spots were detected using anti-HuR antibody. In control cells, two spots were observed for HuR, one at the acidic end towards pH 3 and the other at the basic end towards pH 10. The spots had the same molecular weight, suggesting the presence of two sub-populations of HuR with separate charged modifications that did not alter overall protein weight. An additional spot, slightly more acidic than the one at pH 10 (marked with an asterisk in Fig 1A), was observed upon HCV RNA transfection (Fig 1A, B), indicating a negatively charged modification on HuR.

**Figure 1.**
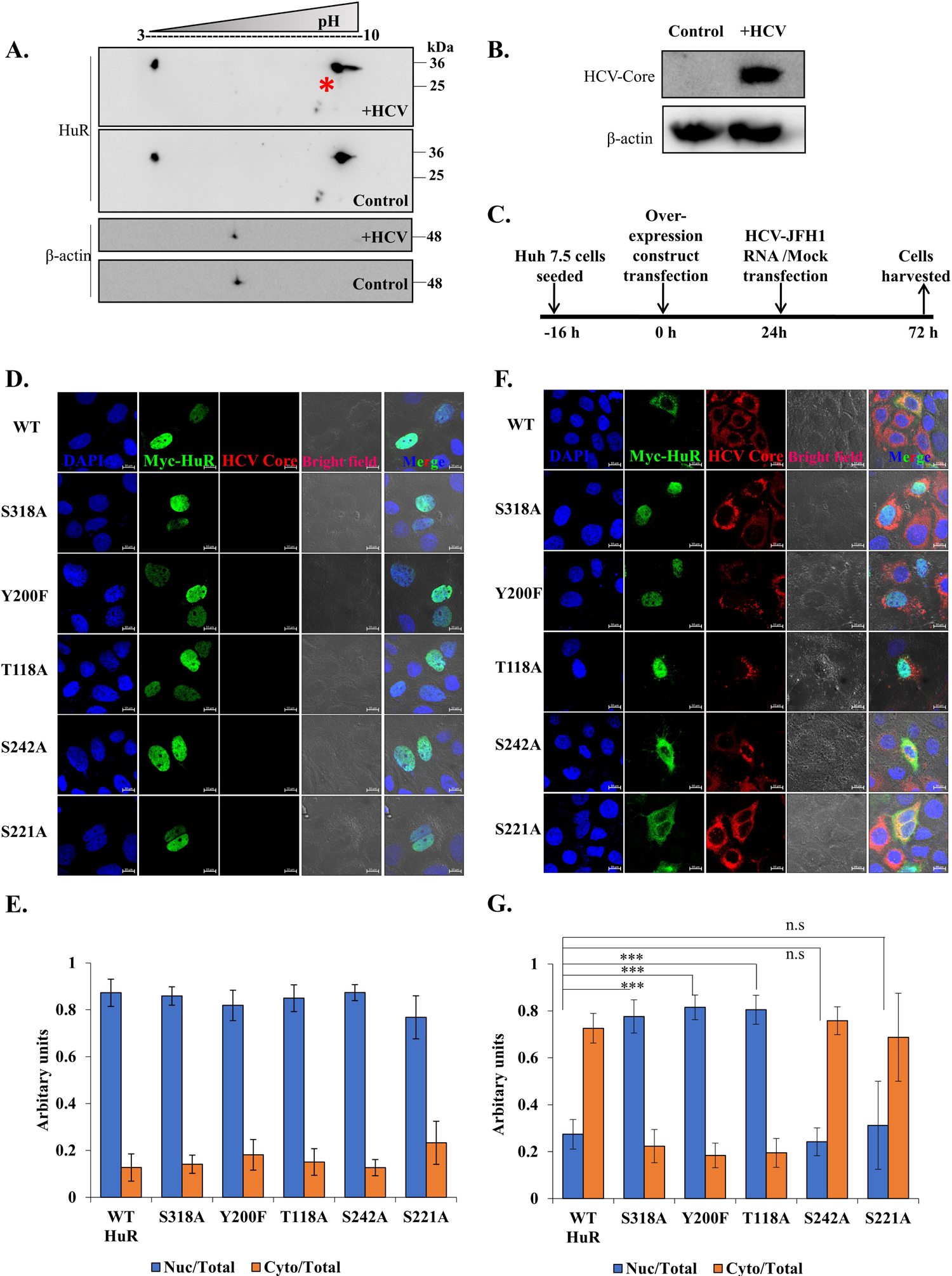
HuR gets phosphorylated upon HCV infection. **(A)** Huh7.5 cells were transfected with HCV-JFH1 RNA. Mock and transfected cells were harvested after 48h and cell lysates were precipitated using TCA precipitation. The precipitated proteins were run in 2 dimensions as described in the methods, followed by Western blotting of the gel. Western blotting was done with anti-HuR and anti-Actin antibodies. Appropriate HRP-conjugate secondary antibodies were used. **(B)** HCV infection was detected by western blotting using Anti-HCV core antibody for same lysates run on 12% SDS-PAGE. **(C)** Schematic for experimental protocol followed. **(D)** Huh 7.5 cells were transfected with Myc-tagged overexpression construct of WT HuR, S318A HuR mutant, S242A HuR mutant, Y200F HuR mutant, T118A HuR mutant and S221A HuR mutant and mock transfected. Cells were processed for immunofluorescence staining using Alexa Fluor conjugated secondary antibody against Myc (Green) and viral protein Core (Red). The nucleus was stained with DAPI (Blue). Scale bar represents 10 µm. **(E)** Nuclear and cytoplasmic ratio for overexpressed HuR (Myc) was quantified for images in (D) using Zen 2.3 lite software. **(F)** Images of HCV transfected cells following the schematic in (C). **(G)** Nuclear and cytoplasmic ratio for overexpressed HuR (Myc) was quantified for images in (F) using Zen 2.3 lite software. n=10. Student t-test was performed for statistical analysis. *= p<0.05, **= p<0.01, ***=p<0.001.

Phosphorylation is a post-translational modification that imparts a negative charge on proteins. Therefore, we examined the involvement of phosphorylation on HuR following HCV infection. Some phosphorylation sites known to impact HuR subcellular localization include T118, Y200, S221, S242, and S318. The phospho-dead mutants of these sites were generated using site-directed mutagenesis, wherein the serine or threonine residues were replaced by alanine (T118A, S221A, S242A, S318A), and tyrosine was replaced by phenylalanine (Y200F). The mutants were verified by sequencing and employed in the relocalization assay. Endogenous HuR translocates from the nucleus to the cytoplasm 48h after HCV-RNA transfection (9). If phosphorylation of a particular residue is crucial for guiding this translocation, the corresponding phospho-dead mutant would prevent the cytoplasmic relocalization of the mutant protein upon viral infection. Wild-type (WT) HuR overexpression was used as a positive control for HuR localization. Using immunostaining, the subcellular localization of WT and mutant HuR proteins was detected in uninfected control and HCV-infected cells. Myc-tagged WT and mutant HuR was overexpressed in Huh7.5 cells, followed by HCV-RNA transfection (Fig 1C). An anti-Myc-tag antibody was used to visualize the overexpressed protein, the HCV Core protein was used to identify HCV-infected cells, and DAPI was used to mark the nuclei. In untransfected control cells, WT and all mutant HuR proteins localized to the nucleus (Fig 1D, E). HCV-RNA transfection led to the relocalization of WT HuR to the cytoplasm. The same was observed for HuR mutants S221A and S242A. In contrast, mutants S318A, Y200F, and T118A remained localized to the nucleus even after 48h of HCV-RNA transfection (Fig 1F, G). This finding suggests that phosphorylation of S318, Y200, and T118 might be involved in relocalization of HuR to the cytoplasm upon HCV infection. The relocalisation of WT and S318A HuR was also analysed by nuclear-cytoplasmic fractionation, wherein WT/S318A HuR expressing Huh7.5 cells were either Mock transfected or transfected with HCV-RNA and localisation of HuR examined by western blotting using anti-HuR antibody (Fig S1C, D). We observed that the ratio of Nuclear to cytoplasmic intensity of overexpressed WT HuR decreased, but that of S318A HuR slightly increased upon HCV-RNA transfection, suggesting and corroborating the imaging results of nuclear retention of S318A HuR upon HCV RNA transfection. The effect on relocalisation of WT and mutant HuR was marginal because the percentage of cells co-transfected with the HuR overexpression construct and HCV-RNA is very low and therefore, immunofluorescence staining serves as a better technique for monitoring the relocalisation.

### HuR is differentially phosphorylated upon HCV infection

To validate HCV induced HuR phosphorylation, total phosphorylated proteins were immunoprecipitated using pan-anti-phospho-Ser/Thr and pan-anti-phospho-Tyr antibodies. The presence of HuR in the pull-down fraction of both antibodies confirmed HuR phosphorylation in cells (Fig 2A, B). HCV core exhibited slight phosphorylation in both Ser/Thr and Tyr pull down. HCV Core protein has Ser, Thr and Tyr motifs which are known and predicted to be phosphorylated. One of the Tyrosine motifs is conserved in all the HCV genotypes and is also reported to be essential for HCV assembly (Tyr 136) (38). There is also tyrosine phosphorylation site prediction in HCV Core (Tyr 86) (39). To identify the phosphorylation sites on HuR, the latter was immunoprecipitated from control or HCV-RNA-transfected cells, and the pull-down fraction was subjected to mass-spectrometric analysis (Fig 2C, D). The results revealed six previously characterized sites, including T80, S88, T143, S146, S202, and S318, as well as two new sites, S135 and T293 (Fig 2E). The abundance of phosphorylated peptides at a particular site was calculated as the fraction of phosphorylated peptides to total peptides observed for that site. Quantification was performed for S318, which was previously found to be regulated by HCV infection, as well as for S202, which is known to regulate HuR localization, and S135, which has not yet been characterized. The percentage of phosphorylated S318 increased in all three sets of HCV-RNA transfected cells compared to control cells (Fig 2F). S202 phosphorylation varied among separate sets of experiments (Fig 2G); whereas S135 phosphorylation was reduced upon HCV-RNA transfection in all three sets (Fig 2H). Differential phosphorylation at distinct HuR sites might explain the absence of an overall variation (Fig 2A, B). Because S318 phosphorylation was higher following HCV RNA-transfection and was found to be important for HuR relocalization (Fig 1F, G), this site was selected for further studies.

**Figure 2.**
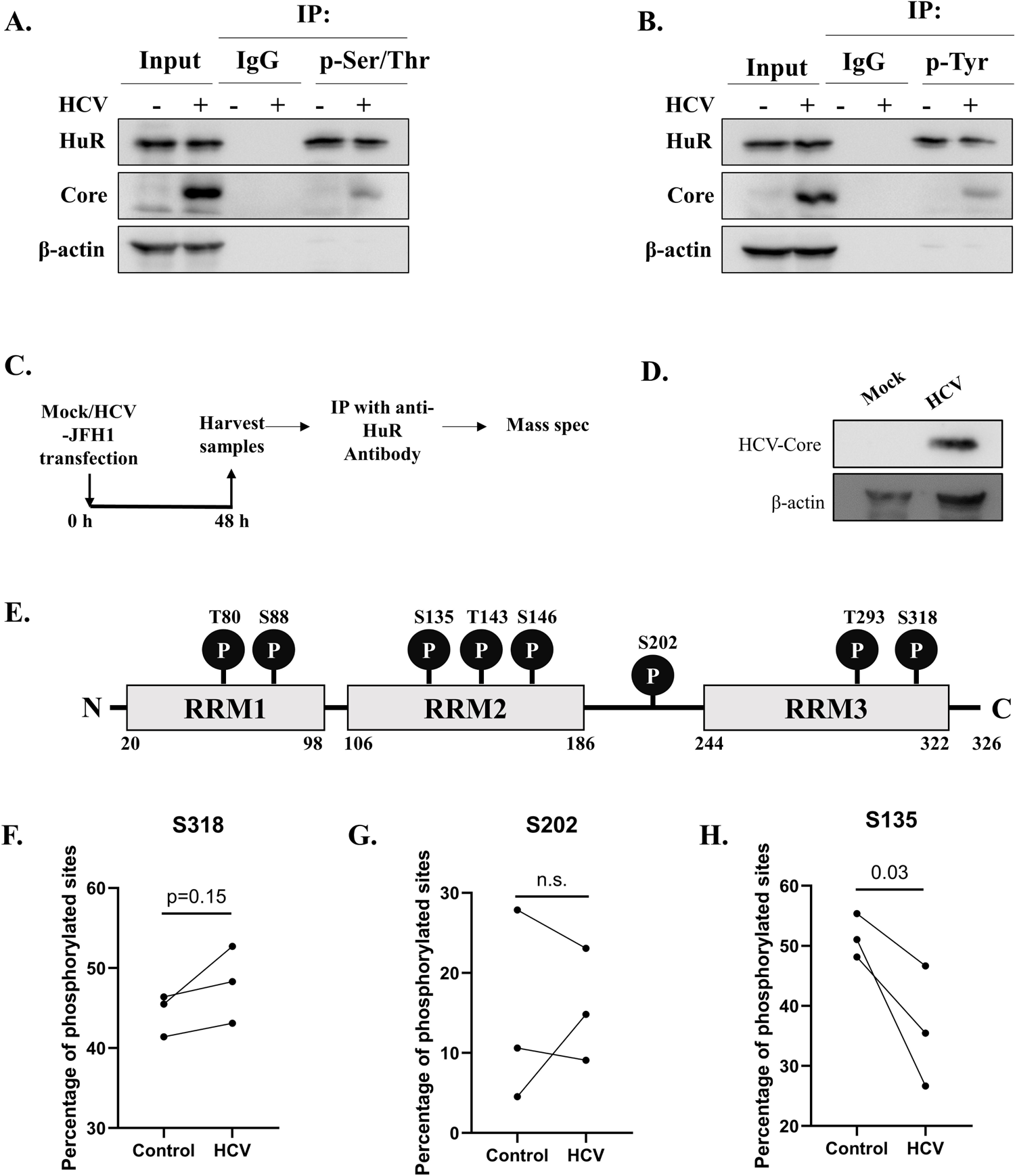
Differential phosphorylation on HuR upon HCV infection. **(A, B)** Huh7.5 cells were mock transfected or transfected with HCV-JFH1 RNA and after 48h, immunoprecipitation was performed using either anti-p-Ser/Thr antibody (A) or anti-p-Tyr antibody (B). Associated HuR was detected by western blot. Western blot for Core was done for confirmation of HCV infection and β-actin was used as loading control. **(C)** Identification of phosphorylation sites on HuR. Immunoprecipitation of HuR was performed from mock or HCV-JFH1 RNA transfected cells and the phosphorylation sites detected using mass-spectrometry. **(D)** HCV infection was confirmed by western blotting for Core using anti-core antibody. **(E)** The phosphorylation sites obtained in mass-spectrometry data were mapped on HuR schematic diagram. **(F-H)** Percentage of phosphorylated S318 (F), S202 (G), S135 (H) sites were calculated as the ratio of peptides containing phosphorylation to the total number of peptides containing respective site. Each dot pair represents a separate set of experiment.

### Phosphorylation of S318 influences HCV replication

The location of the S318 site on HuR was visualized using Pymol software. The X-ray structure of HuR RRM3 (PDB ID: 6GD2) complexed with RNA indicated that the S318 site was on the surface of the protein and interacted directly with the bound RNA (Fig 3A). Accordingly, phosphorylation of S318 might affect the RNA binding of HuR. To study the impact of phosphorylation on the affinity for HCV-RNA, we used surface plasmon resonance. Biotinylated HCV 3’-UTR RNA was immobilized on a streptavidin-coated SPR chip (SA Chip) and recombinant purified WT, S318A, and S318D HuR proteins were passed over as analytes to calculate the binding affinity (Fig 3B–D). The Kd values for WT and S318A HuR were 25.7 nM and 46.2 nM, respectively; whereas the phospho-mimic mutant S318D showed approximately five-fold better affinity (Kd 6.36 nM) (Fig 3E). This result indicates that, along with relocalization to the cytoplasm, S318 phosphorylation of HuR also augments the binding affinity for HCV 3’-UTR. To assess the impact of S318 phosphorylation on HCV replication, WT, S318A, and S318D constructs were overexpressed in Huh7.5 cells, followed by HCV-RNA transfection (Fig 3F). The effect on replication was checked by analyzing the levels of HCV negative-strand RNA compared to the vector-overexpression control. Viral replication was increased upon overexpression of WT HuR but not S318A HuR, which did not relocalize to the cytoplasm and was rescued by overexpression of S318D HuR (Fig 3G, H). We analysed the S318D distribution in untreated cells and observed increased but not complete cytoplasmic localization as compared to WT HuR (Fig S2). These findings confirm the importance of S318 phosphorylation in HuR localization and hence, HCV replication.

**Figure 3.**
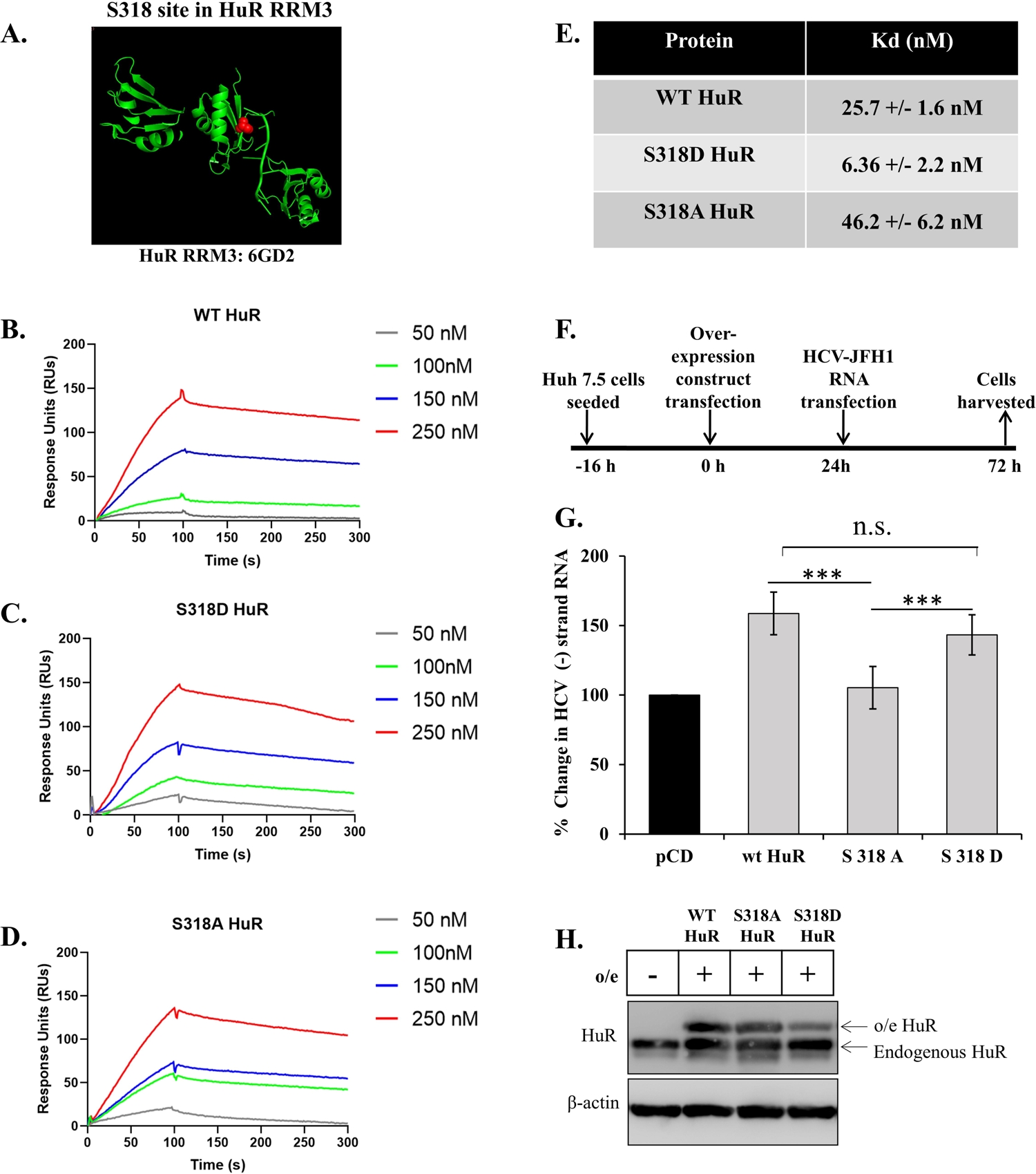
S318 site phosphorylation influences HCV-RNA replication. **(A)** S318 site in HuR RRM3 in association with AU-rich RNA. The red balls indicate S318 site, directly involved in interaction with associated RNA. **(B-D)** Sensorgrams for interaction of (B) WT, (C) S318D HuR and (D) S318A HuR with HCV 3’UTR at indicated concentrations. The y-axis represents the change in Response Units in Association and dissociation phases and x-axis represents time. **(E)** Table for the average *Kd* value obtained for HuR binding to HCV 3’UTR. *Kd* values were calculated as *Kd* =*k*off/*k*on where *k*on represent rate constant of association phase of 100s and *k*off represents rate constant of dissociation phase of 200s using all the 4 concentrations graphs in each protein (B-D) and the average was calculated. **(F)** Schematic for the workflow. **(G)** Following the workflow in (F), total cellular RNA was isolated, and HCV negative strand RNA detected by real time PCR (n=3). **(H)** Western blotting for HuR was performed following the workflow in (F). o/e denotes the overexpression of described WT/mutant HuR. Student t-test was performed for statistical analysis. *= p<0.05, **= p<0.01, ***=p<0.001.

### PKC-**δ** activity is required for HCV-mediated cytoplasmic export of HuR

Once the involvement of S318 phosphorylation in guiding HuR localization was ascertained, we determined the role of the kinase responsible for this modification. An earlier study suggested that PKC-δ participated in the phosphorylation of S318 on HuR and its nucleocytoplasmic shuttling (17, 26). Therefore, the involvement of PKC-δ in regulating HuR localization upon HCV infection was investigated. The levels of total and phosphorylated PKC-δ at different time points after HCV RNA transfection were analyzed. An increase in p-PKC-δ/total PKC-δ after 48h of HCV-RNA transfection was observed, suggesting the activation of PKC-δ upon HCV RNA transfection (Fig 4A). We also observed an increase in cleavage of PKC-δ upon HCV RNA transfection (Fig S3). This cleavage product retains the catalytic subunit of PKC-δ, while freeing it from the regulatory subunit and is known to be transported to the nucleus (40). This cleaved subunit might initiate the cascade to HuR phosphorylation for its cytoplasmic export. To visualize the interaction between HuR and PKC-δ in cells, immunoprecipitation of HuR was performed in mock (control) and HCV-virus infected cells, and the presence of PKC-δ in the pull-down fraction confirmed their physical interaction (Fig 4B). The increase in HuR associated with PKC-δ indicated a stronger interaction in cells upon HCV infection. The association was further strengthened by reverse pull-down, wherein immunoprecipitation of PKC-δ was performed and increased HuR association was observed upon HCV infection (Fig 4C). To correlate this finding with HuR localization, an inhibitor of PKC-δ (rottlerin) was used. The effect of PKC-δ inhibition on HCV-induced HuR relocalization was assessed. Huh7.5 cells were transfected with HCV-JFH1 RNA and 3 µM rottlerin was added to the medium after 6h of transfection (Fig 4D). The cells were incubated with the inhibitor for 48h, after which HuR localization was visualized by confocal microscopy. In the absence of the inhibitor, HuR relocated from the nucleus to the cytoplasm upon HCV RNA transfection (Fig 4E, F). The presence of the inhibitor did not alter the localization of HuR in control cells, but it prevented its relocalization to the cytoplasm upon HCV RNA transfection. These results confirmed the involvement of PKC-δ in altering the localization of HuR upon HCV infection.

**Figure 4.**
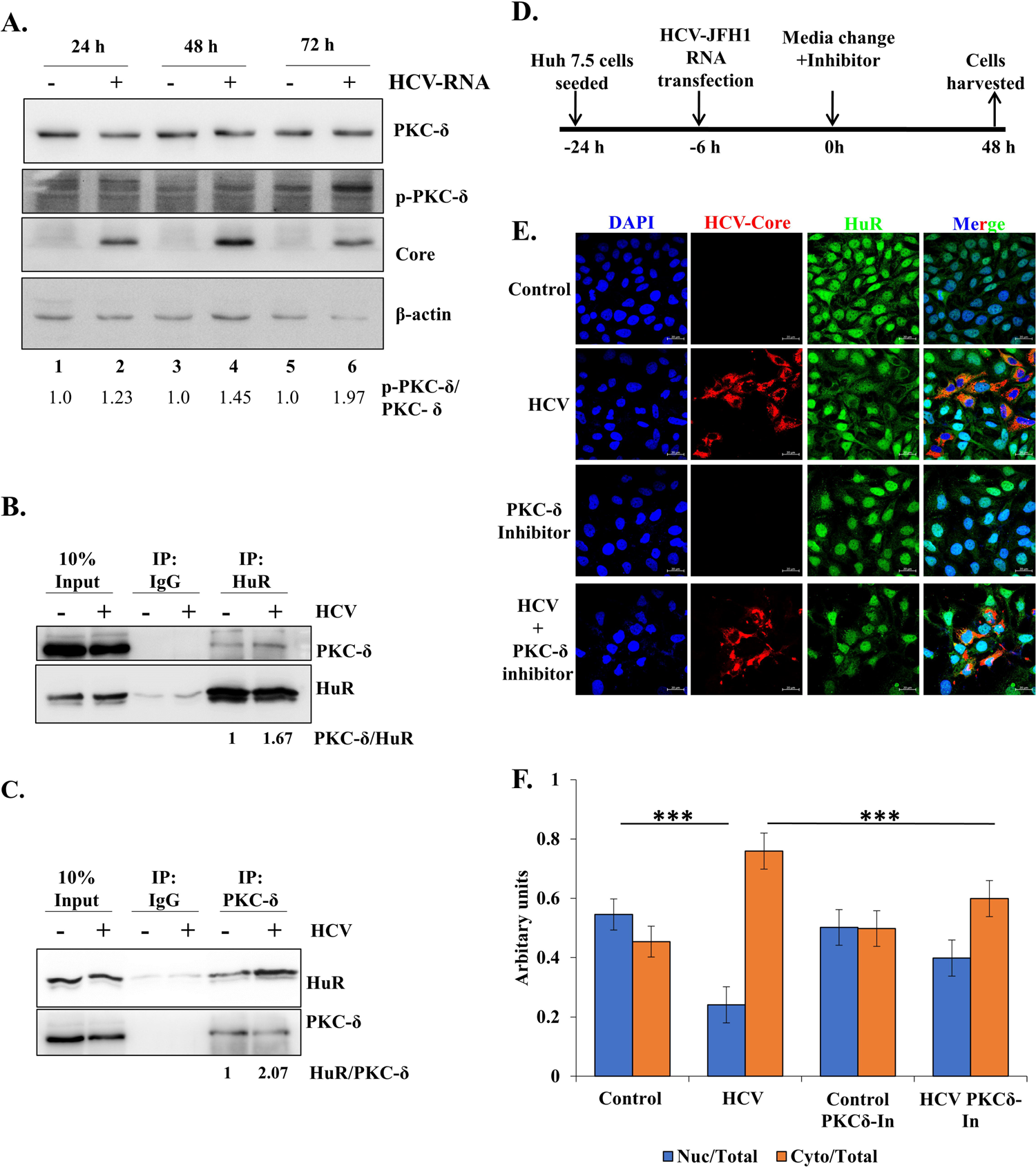
Involvement of PKC-δ in HuR localisation. **(A)** Huh7.5 cells were transfected with HCV-JFH1 RNA and cells harvested at indicated time-points. Western blotting was done with anti-PKCδ, anti-p-PKCδ, anti-Core and anti-actin antibodies. Appropriate HRP-conjugate secondary antibodies were used. **(B)** Immuno-pulldown of HuR was performed from Huh7.5 cells, post 48h of HCV-JFH1 virus infection and associated PKC-δ was checked by western blotting with specific antibody. The values at bottom represent ratio of densitometry values of PKC-δ in the IP fraction to that of HuR in IP fraction. **(C)** Immuno-pulldown of PKC-δ was performed from Huh7.5 cells, post 48h of HCV-JFH1 virus infection and associated HuR was checked by western blot using HuR-specific antibody. The values at bottom represent ratio of densitometry values of HuR in the IP fraction to that of PKC-δ in IP fraction. **(D, E)** Huh7.5 cells were treated with infectious HCV-JFH1 virus in the presence of 3µM of Rottlerin and immunofluorescence staining was carried out at 24h post infection using Alexa Fluor conjugated secondary antibodies against HCV-NS3 (Red) or HuR (Green). The nucleus was counterstained with DAPI. Scale bar represents 10 µm. **(F)** Nuclear and cytoplasmic ratio of HuR (green) was quantified for images in (F) using Zen 2.3 lite software. n=30. Student t-test was performed for statistical analysis. *= p<0.05, **= p<0.01, ***=p<0.001.

### PKC-**δ** regulates the HCV life cycle through HuR

Given that HuR is involved in viral RNA replication, we investigated the effect of rottlerin on HCV-RNA levels. Huh7.5 cells were transfected with HCV-JFH1 RNA and 3 µM rottlerin was added to the medium for 24h (Fig 5A). After 24h, the abundance of HCV-negative strand RNA in the cells was quantified by real-time PCR, which showed that addition of rottlerin decreased HCV-RNA levels by > 90% (Fig 5B). Similarly, confocal microscopy indicated that the percentage of HCV positive cells dropped from ∼20% in untreated cells to <2% in rottlerin-treated cells (Fig 5C, D). In all assays, the PKA inhibitor KT5720 was used as a negative control, which did not show any effect on HuR localization upon HCV infection. However, we observed an increase in replication upon PKA inhibitor treatment which could be because of the regulation of localisation of another negative regulator of HCV replication, PTB (41). The effect on viral replication was also assayed using siRNA mediated knockdown of PKC-δ. Huh7.5 cells were transfected with HCV-JFH1 RNA after 24h of siRNA transfection. 48h post HCV RNA transfection, HCV-negative strand RNA was quantified. We observed a dose-dependent decrease in HCV-negative strand RNA with increasing concentration of siRNA targeting PKC-δ (Fig 5 E, F).

**Figure 5.**
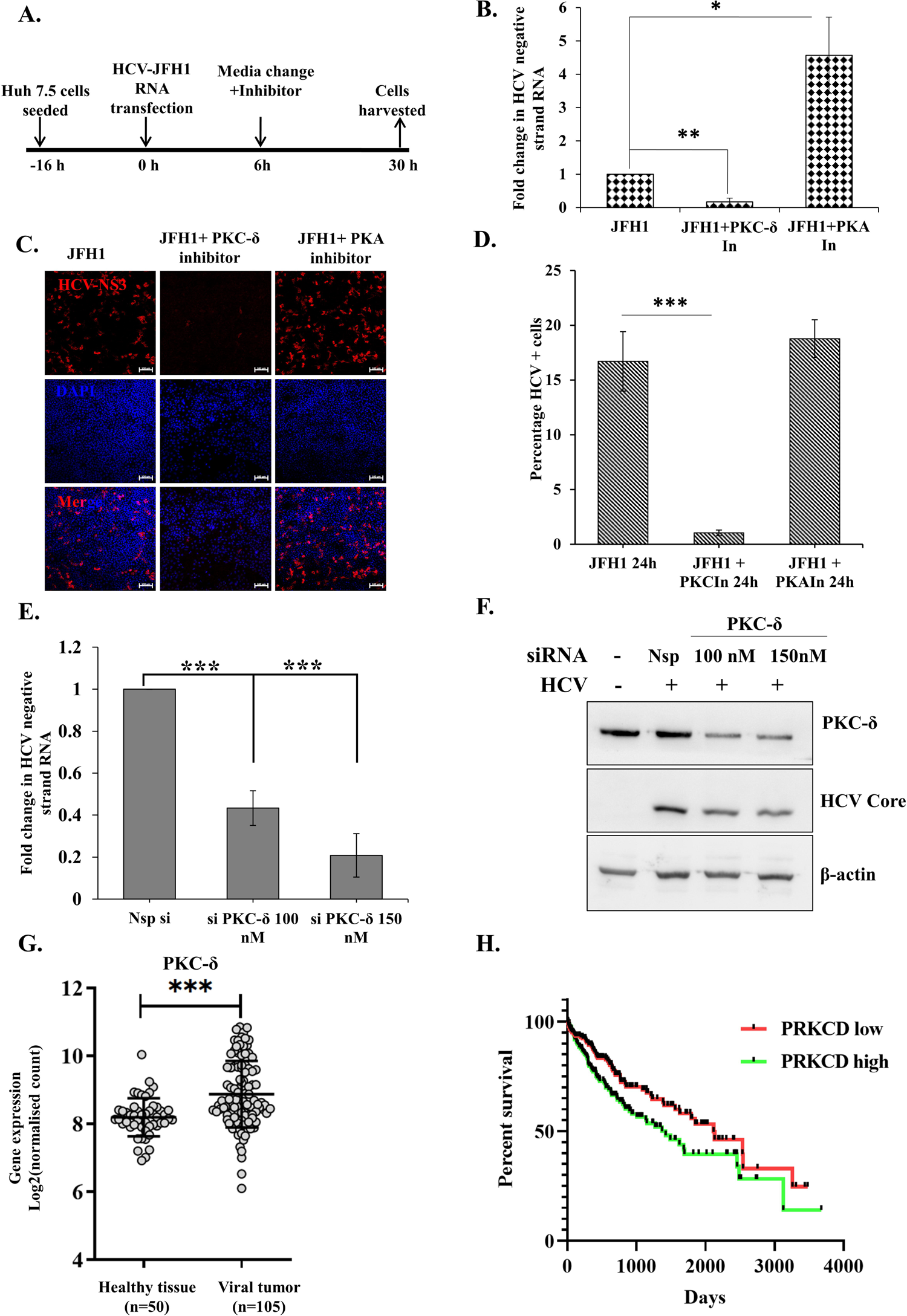
Role of PKC-δ. In HCV life cycle. **(A)** Workflow of the experiment. **(B)** According to the workflow, cells were harvested after 24h of addition of inhibitors. Total RNA was isolated from the cells and HCV negative strand RNA was detected using RT-qPCR. Fold change in RNA levels was calculated as compared to RNA levels in cells with no inhibitor treatment. (n=3) **(C)** Huh7.5 cells were transfected with HCV-JFH1 RNA in the presence of 3µM of Rottlerin or 10μM KT5720 and immunofluorescence staining was carried out at 24h post infection using Alexa Fluor conjugated secondary antibodies against HCV-NS3 (Red). The nucleus was counterstained with DAPI. Scale bar represents 100µm. **(D)** Quantification of percentage of HCV positive cells in the confocal images in panel C. (n=3) **(E)** Huh7.5 cells were transfected with either non-specific siRNA (Nsp si) or indicated concentrations of siRNA targeting PKC-δ (si PKC-δ). 16h post transfection, cells were transfected with HCV-JFH1 RNA, and 48h post JFH1-RNA transfection cells were harvested, and HCV negative strand RNA levels determined by qRT-PCR using HCV specific primers. **(F)** The extent of PKC-δ silencing in (E) was determined by western blotting by using anti-PKC-δ antibody. HCV-Core protein served as the marker for HCV-RNA transfection. β-actin was used as loading control. **(G)** Expression level of PKC-δ (PRKCD gene) in healthy tissue and Virus induced primary HCC tissue samples from TCGA database. **(H)** Kaplan Meier plot for survival probability for Virus induced primary HCC patients with varying level of expression of PKC-δ. Data taken from TCGA database. Student t-test was performed for statistical analysis. *= p<0.05, **= p<0.01, ***=p<0.001.

The expression of PKC-δ in healthy and hepatocellular carcinoma patients (Virus induced HCC patient positive for either HCV, HBV or both) in The Cancer Genome Atlas (TCGA) database was analyzed. Increased PKC-δ expression in virus-induced hepatocellular carcinoma patients suggested its involvement in disease progression (Fig 5G), and survival probability was inversely proportional to PKC-δ expression levels (Fig 5H). This result suggested that patients with elevated PKC-δ would have more and early relocalisation of HuR upon HCV infection, which would lead to enhanced viral replication and, hence, increased likelihood of severe disease progression.

### HCV non-structural proteins increase the cytoplasmic abundance of HuR

Activation of PKC-δ and relocalization of HuR to cytoplasm seemed to be a viral strategy for its efficient replication. Therefore, candidate viral protein(s) involved in PKC-δ activation were investigated. Over-expression of either the structural protein, Core or all the non-structural proteins together through pSGR-Luc construct was performed in Huh7.5 cells and localization of HuR was assessed after 48h of overexpression. Viral non-structural proteins were found to be sufficient to cause the relocalization of HuR (Fig 6A, B). Among non-structural proteins, NS3 is a major pathogenic protein that interacts with multiple host proteins and kinases (42–45). To assess the physical interaction between NS3 and PKC-δ, interaction studies for NS3 and PKC-δ were performed in Huh7.5 cells. Myc-tagged NS3 overexpression construct was transfected in Huh7.5 cells and 48h post transfection, PKC-δ was pulled down from the cell lysate to assess co-immunoprecipitation of myc-tagged NS3. PKC-δ pull-down could immunoprecipitated NS3 from the cell lysate, establishing their physical association (Fig 6C). This interaction was further confirmed by the colocalization of NS3 and PKC-δ in HCV-JFH1 RNA transfected cells with a colocalization coefficient of 0.898 (Fig 6D). The colocalization assay was performed using HCV infection as well. The staining for HCV-NS3 and HCV-Core was performed to examine their interaction with PKC-δ (Fig 6E). This yielded a colocalization coefficient of 0.184 for Core and PKC-δ colocalization and a colocalization coefficient of 0.85 for NS3 and PKC-δ colocalization, suggesting the interaction between PKC-δ and NS3 and not Core in HCV infected cells. These assays provide the evidence of direct interaction of HCV-NS3 protein with PKC-δ, which could activate it for HuR phosphorylation at S318 and hence its cytoplasmic localisation upon HCV infection.

**Figure 6.**
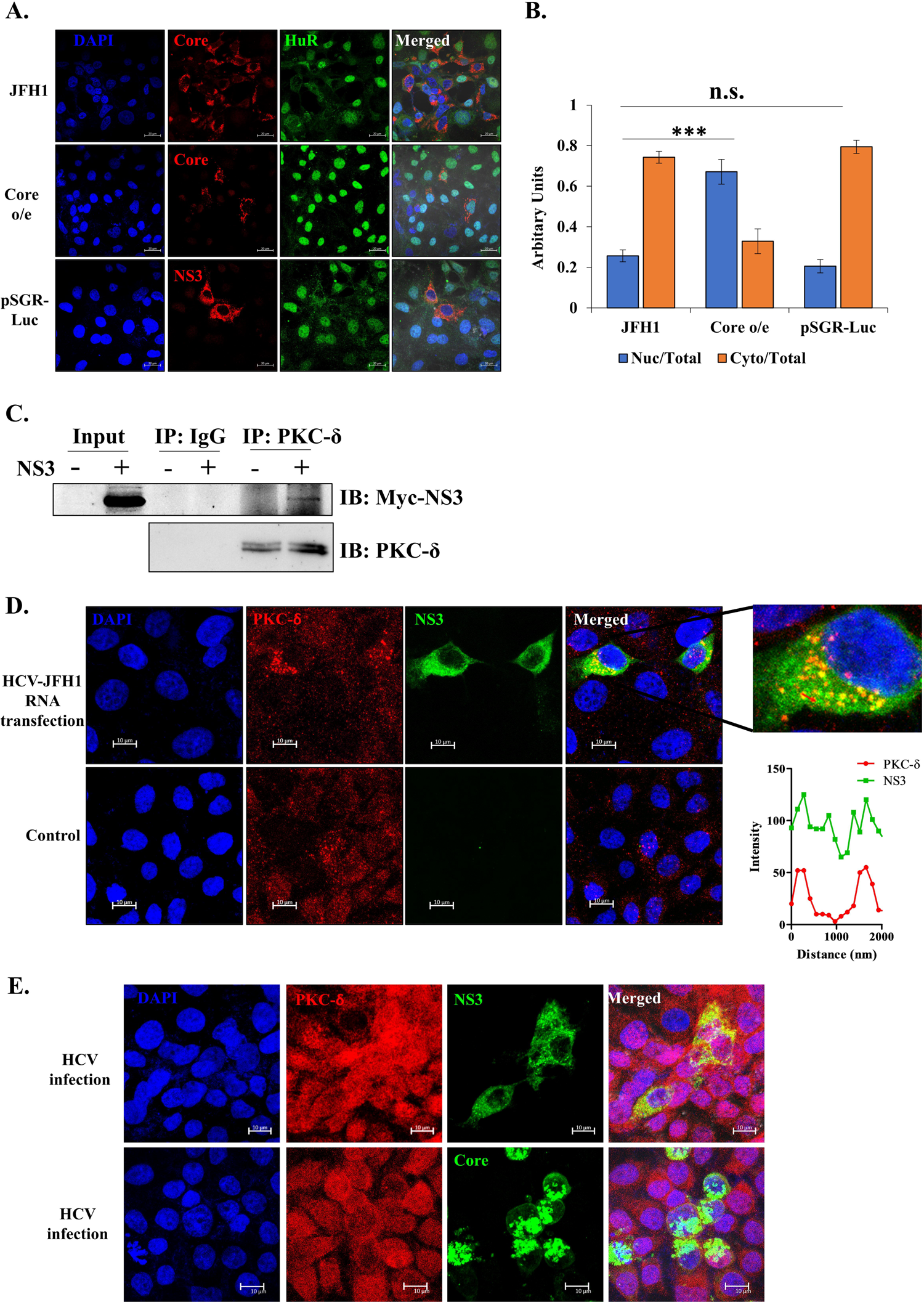
Viral protein involved in relocalisation of HuR. **(A)** Huh7.5 cells were transfected with plasmid expressing HCV protein as indicated. Immunofluorescence staining was carried out at 48h post transfection using Alexa Fluor conjugated secondary antibodies against HuR (Green) and indicated viral proteins (Red). The nucleus was counterstained with DAPI. Scale bar represents 20µm. **(B)** Nuclear and cytoplasmic ratio for HuR was quantified for images in (A) using Zen 2.3 lite software. n=30. Student t-test was performed for statistical analysis. *= p<0.05, **= p<0.01, ***=p<0.001. **(C)** Myc-tagged NS3 was overexpressed in Huh7.5 cells and 48h post transfection, cells were harvested for coimmunoprecipitation. Anti-PKC-δ antibody was used to pull down PKC-δ from vector control (pCDNA3.1) or Myc-NS3 transfected cell lysates. Western blotting was performed using anti-Myc antibody to detect the presence of NS3 in pull-down fraction. **(D)** Huh7.5 cells were transfected with HCV-JFH1 RNA. Immunofluorescence staining was carried out at 48h post transfection using Alexa Fluor conjugated secondary antibodies against NS3 (Green) and PKC-δ (Red). The nucleus was counterstained with DAPI. Scale bar represents 10µm. An enlarged image of infected cell and the line profile for PKC-δ and NS3 intensity over the indicated arrow in the enlarged image is depicted in the inset. Colocalization coefficient was calculated using Zeiss software. **(E)** Huh7.5 cells were infected with HCV-JFH1 virus. Immunofluorescence staining was carried out at 48h post transfection using Alexa Fluor conjugated secondary antibodies against NS3, Core (Green) and PKC-δ (Red). The nucleus was counterstained with DAPI. Scale bar represents 10µm. Colocalization coefficient was calculated using Zeiss software.

### AMPK is involved in cytoplasmic retention of HuR upon HCV infection

Viral infection leads to HuR relocalization to the cytoplasm; however, at the same time, it is essential for the virus that HuR is prevented from reentry in the nucleus. The nuclear localization signal for HuR is not very well characterized. AMPK has been shown to phosphorylate and acetylate importin-α1, which carries newly synthesized HuR to the nucleus. Here, we assessed whether this mechanism was disrupted during HCV infection, thus allowing HuR to be retained in the cytoplasm. Huh7.5 cells were treated with an AMPK inhibitor (Compound C) or an AMPK activator (A769662) following HCV-RNA transfection. HuR localization was assessed by confocal microscopy 48h after drug treatment. Inhibition of AMPK activity did not alter HuR localization, whereas activation of AMPK led to increase in nuclear and decrease in cytoplasmic HuR and prevented complete cytoplasmic relocalisation of HuR, even upon HCV RNA transfection (Fig 7A, B). NS5A overexpression alone could not cause HuR relocalisation to cytoplasm (Fig. S4), suggesting that the trigger of NS3 mediated S318 phosphorylation could be required for initial HuR transport to the cytoplasm.

**Figure 7.**
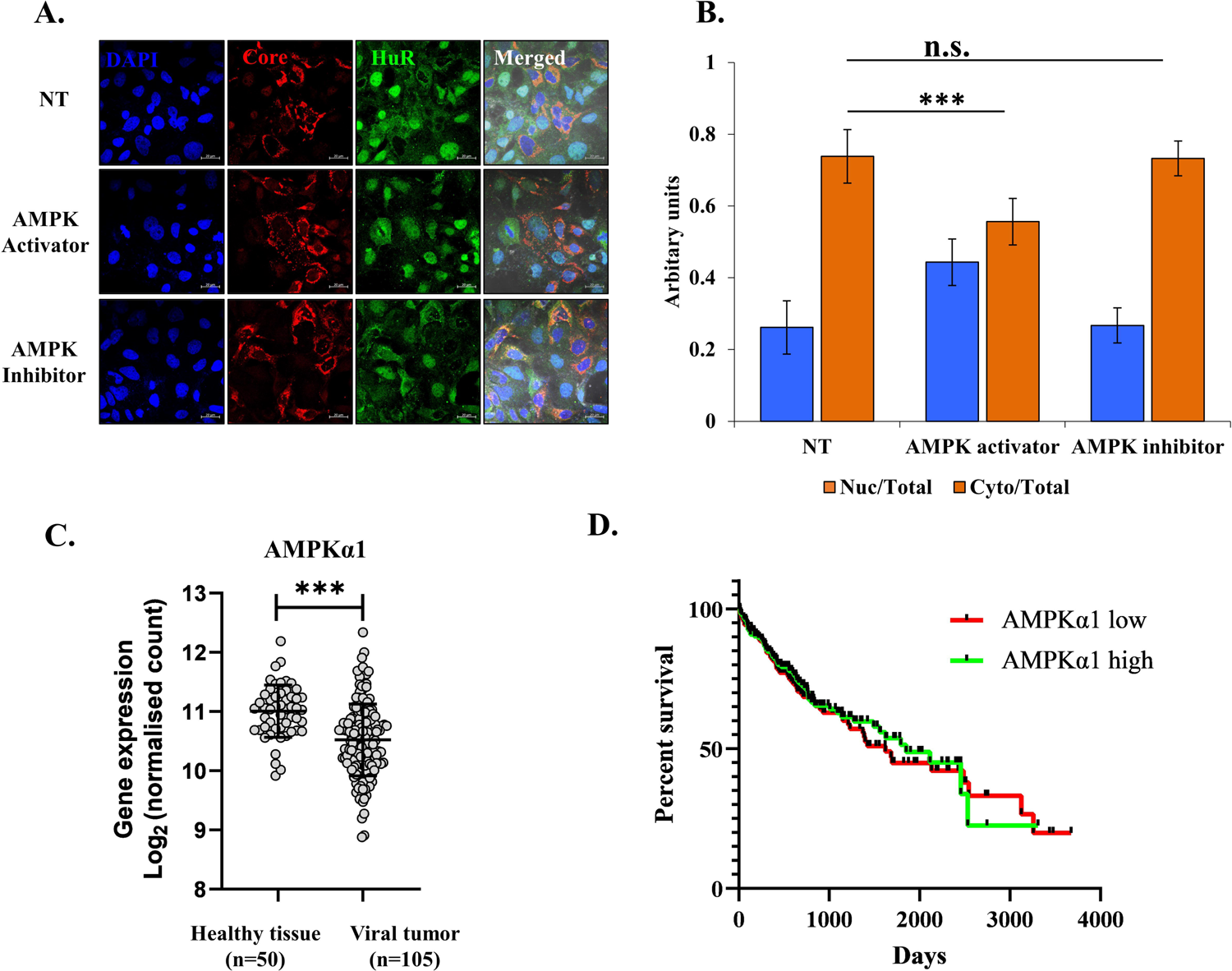
AMPK in relocalisation of HuR. **(A)** Huh7.5 cells were transfected with HCV-JFH1 RNA in the presence of 10µM of AMPK inhibitor Compound C (CC) or 100mM A76 (AMPK activator) and immunofluorescence staining was carried out after 48h using Alexa Fluor conjugated secondary antibodies against HCV-Core (Red) and HuR (Green). The nucleus was counterstained with DAPI. Scale bar represents 20µm. **(B)** Nuclear and cytoplasmic ratio of HuR was quantified for images in (A) using Zen 2.3 lite software. n=30. **(C)** Expression level of PRKAA2 (AMPKα2) in healthy tissue, HCC primary tumor samples and Virus induced primary HCC tissue samples from TCGA database. **(D)** Kaplan Meier plot for survival probability for Virus induced primary HCC patients with varying level of expression of AMPKα2. Data taken from TCGA database. Student t-test was performed for statistical analysis. *= p<0.05, **= p<0.01, ***=p<0.001.

To correlate AMPK activity with hepatocellular carcinoma progression, TCGA database AMPK levels were analyzed. A decrease in total AMPKα1 levels was observed upon virus-induced tumor progression (Fig 7C). No significant correlation was observed between survival probability and AMPK-α1 levels (Fig 7D). The viral non-structural protein NS5A inhibits AMPK activity (46, 47), and we showed here that AMPK inhibition could prevent complete cytoplasmic relocalisation of HuR upon HCV infection.

Our results uncovered the participation of multiple viral proteins in achieving increased cytoplasmic abundance of HuR. We describe the underlying mechanism wherein viral RNA translates upon entry into the cell, producing viral proteins. Viral NS3 protein activates PKC-δ, which phosphorylates HuR on S318, leading to its relocalization to the cytoplasm, where it is available for the replication of cytoplasmic viral RNA. At the same time, viral NS5A protein inhibits AMPK, which retains relocalized and newly synthesized HuR in the cytoplasm (Fig 8). The proposed mechanism depicts a coordinated effort to regulate a host RNA binding protein, so it can be exploited to guide the viral life cycle.

**Figure 8.**
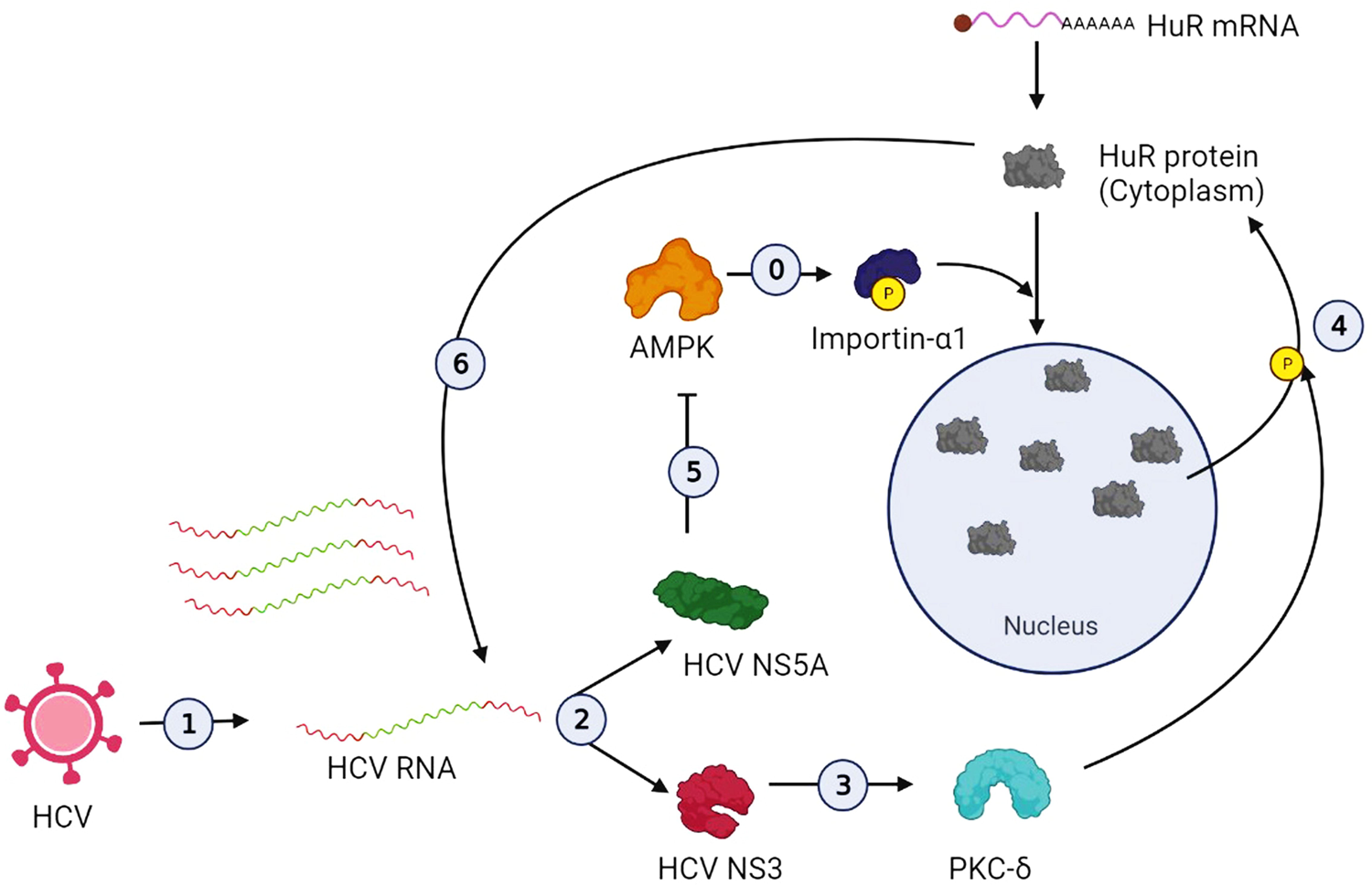
Proposed mechanism of HuR relocalisation upon HCV infection. (0) Cellular mechanism of HuR transport to nucleus. AMPK phosphorylates Importin-α1 which guides HuR to the nucleus. (1) HCV entry in the cells followed by Viral RNA release in the cell cytoplasm. (2) Viral RNA translates to produce structural and non-structural proteins. (3) Non-structural protein 3 (NS3) activates PKC-δ. (4) PKC-δ phosphorylates HuR on S318 residue, guiding its cytoplasmic export. (5) Non-structural protein 5A (NS5A) inhibits AMPK to retain the cytoplasmically relocalised and new synthesised HuR in the cytoplasm. (6) The cytoplasmically localised HuR assists in viral replication by binding to viral 3’UTR.

## Discussion

HCV is a hepatotropic virus that can cause chronic infection, leading to hepatocellular carcinoma. Host factors play an essential role in regulating the virus life cycle and cellular pathogenesis. HCV is a positive-strand RNA virus, whose RNA is replicated and translated in the cytoplasm. RNA binding proteins (RBPs) such as HuR bind to viral RNA and regulate these processes.

HCV is a completely cytoplasmic virus with no nuclear history of viral RNA. Therefore, it mediates the relocalization of RBPs to the cytoplasm to enable RNA binding. We show that phosphorylation of the S318 residue on HuR is essential for this relocalization. This site lies in RRM3 of HuR, which is critical for its binding to HCV-RNA. Indeed, we demonstrate that phosphorylation of S318 promotes the binding of HuR to HCV-RNA. Altered S318 phosphorylation levels have been detected in various pathologies such as colon cancer (27), suggesting that the same could happen in hepatocellular carcinoma. We report that S318 phosphorylation is mediated by PKC-δ. Inhibition of PKC-δ by lenvatinib has recently been associated with decreased tumorigenicity in Huh7 and Hep3B hepatocellular carcinoma cells (48). One of the underlying mechanisms behind the pro-proliferative effect of PKC-δ could be its regulation of HuR localization in cells, which directly influences the cellular transcripts involved in tumorigenicity. These reports and the evidence presented herein point to the key role of S318 phosphorylation in the transport of HuR to the cytoplasm and in HCV infection.

HuR is not known to be mutated in many cancers. However, its localization and binding to different RNA transcripts affect cancer progression (49). Reduced AMPK activity, which leads to cytoplasmic localization of HuR, is known to stabilize oncogenic molecules that promote progression of the proliferative phenotype. Here, we show that this mechanism is activated also by HCV, pointing to the significant role of HuR cytoplasmic localization in the progression of HCV-mediated hepatocellular carcinoma. HuR targets include invasive and metastatic RNAs (e.g., MMP-9, MTA1, and uPA), angiogenic RNAs (e.g., VEGF-1 and HIF-1), and cell proliferative RNAs (e.g., EGF, cyclin A, cyclin B1, cyclin E, and cyclin D1), through which it can influence hepatocellular carcinoma progression (50, 51). By stabilizing these target RNAs and increasing their protein products, HuR promotes the transformation from non-cancerous to cancerous cells. Hence, HuR has been proposed as a potential target for cancer treatment (52).

AMPK regulates the availability of HuR to the above RNAs in the cytoplasm. AMPK levels and activity are reduced in patients with hepatocellular carcinoma, and low p-AMPK levels correlate with higher tumor occurrence (53, 54). Regulation of pathogenic RNAs by cytoplasmic HuR may be one of the mechanisms underlying this correlation. AMPK is a double-edged sword, which influences cell proliferation in either way. A recent report has shown that the effect of AMPK on cell proliferation is a function of nutrient abundance. In cancer cells subjected to chronic nutrient deprivation, AMPK inactivation limits cell proliferation (55). This limitation could be explained by direct activation of mTORC1 and mTORC2. Another possibility is the cytoplasmic retention of HuR, which is known to stabilize and enhance the translation of rictor, thereby activating mTORC2 (56). This could be one of the mechanisms of regulation of viral replication in cells and hepatocellular carcinoma progression.

PKC-δ and AMPK are important cellular kinases that direct cell fate under various conditions. HCV exploits these kinases to regulate replication and pathogenesis through HuR. Preventing HCV induced cytoplasmic relocalization of HuR by either targeted PKC-δ inhibition or AMPK activation or by blocking the interaction of non-structural proteins with these kinases could offer novel therapeutic intervention against hepatocellular carcinoma progression.

## Materials and methods

### Plasmids and constructs

*pJFH1*: Plasmid encoding all HCV proteins. The plasmid was linearized using XbaI to synthesise JFH1 RNA (Obtained from Apath,LLC).

*pSGR-JFH1/Luc*: Bicistronic plasmid where the luciferase expression is driven by HCV-IRES while HCV non-structural proteins are expressed by EMCV IRES. The plasmid was linearised by XbaI to synthesise SGR-JFH1/Luc RNA (A kind gift from Dr. Ralf Bartenschlager, Heidelberg University).

*pcDNA3.1-myc-HuR*: The sequence coding for HuR cloned between EcoRI and XhoI sites of pcDNA3.1-myc vector.

*pcDNA3.1-myc-NS3:* The sequence coding for NS3/4A was cloned between EcoRI and XhoI sites of pcDNA3.1-myc vector.

*HuR mutants:* HuR mutants were generated in the pcDNA3.1-myc-HuR background by site directed mutagenesis and the mutations were confirmed by sequencing.

### Cell Lines and Transfections

Huh7.5 cells were maintained in Dulbecco’s Modified Eagle Medium DMEM (Sigma) with 10 % Fetal Bovine Serum. Lipofectamine 2000 (Invitrogen) was used for JFH1 RNA transfections and TurboFect (Thermo Scientific) was used for DNA transfections according to manufacturer’s protocol. For generation of the infectious virus, HCV-JFH1 RNA was *in vitro* transcribed and transfected into Huh7.5 cells. Supernatant from the transfected cells was concentrated and used for viral infection. Uninfected Huh7.5 cells were used as a mock control.

### HCV virus preparation

Huh7.5 cells electroporated with HCV-JFH1 RNA. 72-96h post electroporation, cell supernatant was collected and viral RNA copy number in supernatant determined using standard graph of known number of RNA molecules. Thereafter MOI of infection was determined after calculating specific infectivity of the virus. All virus experiments were done at an MOI of 0.01.

### Nuclear-cytoplasmic fractionation

Nuclear and cytoplasmic extracts were prepared using the SIGMA CelLytic NuCLEAR Extraction Kit as per manufacturer’s protocol. Briefly, cells were lysed in hypotonic lysis buffer and treated with 1 % IGEPAL as per the manufacturer’s recommendations (NXTRACT, SIGMA) to get cytoplasmic extract. The nuclear pellet was then lysed in the extraction buffer. The protein concentration of each extract was determined by Bradford assay. Equal amounts of protein were resolved on a SDS-10 % PAGE followed by western blot using the desired antibodies.

### 2D-gel electrophoresis

The protein lysates for Mock and HCV transfected samples were precipitated using 10 % TCA and pellets washed with acetone. The protein pellets were resuspended in sample rehydration buffer (8 M urea, 2 % [wt/vol] 3-[(3-cholamidopropyl)-dimethylammonio]-1-propanesulfonate [CHAPS], 15 mM dithiothreitol [DTT], and 0.5 % [vol/vol] IPG buffer [pH 3 to 10]). Isoelectric focusing was performed using immobilized pH gradient (IPG) strips (Bio-Rad). IPG strips with a pH range from 3 to 10 were used. For the first dimension, 500 µg of protein samples in 150 µl of rehydration solution was used to rehydrate the IPG strip (7 cm [length]; pH 3 to 10). The IPG strips were passively rehydrated overnight, and then the proteins were focused for 10,000V. h at 20 °C under mineral oil. After focusing, the strips were incubated for 10 min in 2 ml of equilibrium buffer I (6 M urea, 30 % [wt/vol] glycerol, 2 % [wt/vol] sodium dodecyl sulfate [SDS], and 1 % [wt/vol] DTT in 50 mM Tris-HCl buffer [pH 8.8]), followed by equilibrium buffer II (6 M urea, 30 % [wt/vol] glycerol, 2 % [wt/vol] SDS, and 4 % [wt/vol] iodoacetamide in 375 mM Tris-HCl buffer [pH8.8]). After the equilibration steps, the strips were transferred to a 12 % SDS-polyacrylamide gel electrophoresis (PAGE) gel for the second dimension. Protein spots were visualized by silver staining.

### Immunofluorescence staining

For immunofluorescence staining, ∼0.2*10^6^ Huh7.5 cells were seeded on coverslips in a 12-well plate for 14 h followed by transfection of HCV-JFH1 RNA (1μg). After desired time of infection/ RNA transfection, cells were washed twice with 1X PBS and fixed using 4 % formaldehyde at room temperature for 20 min. After permeabilization by 0.1 % Triton X-100 for 2 min at room temperature, cells were incubated with 3 % BSA at 37 °C for 1 h followed by incubation with the indicated antibody for 2 h at 4 °C and then detected by Alexa-633-conjugated anti-mouse or Alexa-488 conjugated anti-rabbit secondary antibody for 30 min (Invitrogen). Images were taken using Zeiss microscope and image analysis was done using the Zeiss LSM or ZEN software tools.

### Western-blot analysis and antibodies

Protein concentrations of the extracts were assayed by Bradford reagent (Bio-Rad) and equal amounts of cell extracts were separated by SDS-12 % PAGE and transferred onto a nitrocellulose membrane (Sigma). Samples were then analyzed by western blot using the desired antibodies, anti-HCV core antibody (ab2740, Abcam), anti-HuR antibody (3A2, Santa Cruz and 07-1735, Merck Millipore), anti-pan-p-Ser/Thr antibody (AP0893, Abclonal), anti-pan-p-Tyr antibody (AP0905, Abclonal), anti-PKC-δ antibody (A7778, Abclonal), anti-p-PKC-δ-T505 (AP0776) followed by the respective secondary antibodies (horseradish peroxidase-conjugated anti-mouse or anti-rabbit IgG; Sigma). Mouse-monoclonal anti-β-actin-peroxidase antibody (A3854, Sigma) was used as a control for equal loading of total cell extracts. Antibody complexes were detected using the ImmobilonTM Western systems (Millipore). NS3 antibody used for immunostaining and co-immunoprecipitation was a kind gift from Prof. Guangxiang (George) Luo, University of Alabama.

### Immunoprecipitation

Huh7.5 cells were scraped in polysome lysis buffer (100 mM KCl, 5 mM MgCl_2_, 10 mM HEPES pH 7.0, 0.5 % NP-40, 1 mM DTT, 100 U/ml RNasin) and kept on cyclomixer for 1 h, followed by spinning at 16000×g for 15 min at 4 °C. The supernatant obtained was precleared with Protein G Sepharose beads for 1 h at 4 °C. The samples were spun at 1000×g for 2 min to pellet the beads, and the supernatant was removed. Protein G Sepharose beads were incubated with 1 ug of respective antibody overnight at 4 °C in a total volume of 200 μl of polysome lysis buffer and added to the precleared lysates followed by incubation for 3 h with continuous mixing on a rotator device at 4°C. The beads were washed three times with polysome lysis buffer. SDS sample buffer was then added to the beads and boiled to release the immunoprecipitated protein, and the supernatant was electrophoresed on a SDS-12 % PAGE.

### Surface Plasmon Resonance

Surface plasmon resonance spectroscopy was performed using a BIAcore3000 optical biosensor (GE Healthcare Lifescience) to study the binding kinetics of HuR with HCV 3’UTR RNA. Biotin-labelled HCV 3’UTR RNA was immobilized on streptavidin-coated sensor chips (GE Healthcare Lifescience) to a final concentration of 300 Resonance Units (RU)/flow cell. RNA–protein interactions were carried out in a continuous flow of Tris buffer (25 mM Tris (pH 7.5), 100 mM KCl, 7 mM β-mercaptoethanol, and 10% glycerol) at 25 °C at a flow rate of 10 µl/min. Increasing concentrations of HuR protein loaded on the biosensor chip for 100 s (characterized as the association phase), followed by a dissociation phase of 300 s with buffer alone. For normalising background non-specific interaction, a blank surface without any RNA was used for simultaneous injections of the sample during the experiment. BIAevaluation software (version 3.0) was used to determine the on rate, *k*on (M^-1^ s^-1^), and off rate, *k*off (s^-1^), using a 1:1 Langmuir binding model. The binding affinity, *Kd* was determined using the following equation: *Kd* =*k*off/*k*on.

### Immunoprecipitation of HuR followed by Mass spectrometry

Huh7.5 cells were transfected with HCV-JFH1 RNA and harvested after 48h of transfection. Total proteins were extracted in IP lysis buffer (Pierce, Thermo Scientific) containing 1× halt-protease and phosphatase inhibitor cocktail (Thermo Fisher Scientific, USA) with intermediate vortexing followed by mild sonication. The lysate was centrifuged at 14,000 × g for 30 min at 4 °C. The supernatant was quantified by the BCA method (Thermo Fisher Scientific, USA). An equivalent amount (4[mg) of proteins from each condition was incubated with 3μg of anti-HuR antibody for overnight at 4[°C. The immunocomplex was captured by using protein G Sepharose 4 fast flow beads (17-0618-01/ GE Healthcare). The immunoprecipitated complex was washed two times with IP lysis buffer and one time with mili-Q. Bound protein complexes were eluted in 50 µl SDS-PAGE 2X Laemmli sample buffer. The samples were resolved on 12% SDS–PAGE and stained with Coomassie stain (0.1% w/v). The gel was subjected to further in-gel mass spectrometry analysis.

Gel slices corresponding to each lane were excised in individual eppendorf and destained using wash buffer (50 mM ammonium bicarbonate (ABC) and 50% ACN). Slices were hydrated with 10 mM DTT (56°C, 30 min), alkylated using 50 mM IAA (30 min, RT, dark), and further dehydrated using 100% ACN before digestion. The available proteins in-gel pieces were digested with MS grade trypsin (Pierce™ Trypsin Protease MS grade, cat no. 1862743, USA) and incubated at 37 °C for 18 h. Digested peptides were extracted by adding 60% ACN with 0.1% FA and 100% ACN with 0.1% FA, followed by ultra-sonication. The digested peptides were collected, vacuum dried, and reconstituted in 40 μl of solvent A (2% (v/v) ACN, 0.1% (v/v) FA in water) and subjected to LC−MS/MS experiments using Sciex 5600^+^ Triple-TOF mass spectrometer coupled with ChromXP reversed-phase 3 μm C18-CL trap column (350 μm[×[0.5[mm, 120 Å, Eksigent, AB Sciex) and nanoViper C18 separation column (75 μm[×[250[mm, 3 μm, 100 Å; Acclaim Pep Map, Thermo Scientific, USA) in Eksigent nanoLC (Ultra 2D plus) system. The binary mobile solvent system was consisted with solvent A (2% (v/v) ACN, 0.1% (v/v) FA in water) and solvent B (98% (v/v) ACN, 0.1% (v/v) FA). The peptides were analysed with 300[nl/min flow rate in a 60 min gradient with a total run time of 75[minutes. The acquisition was executed with conventional data-dependent IDA mode. Each cycle consisted of 250 and 100 ms acquisition time for MS1 (m/z 350−1250 Da) and MS/MS (100–1500 m/z) scans respectively with a total cycle time of 2.8 s. Each fraction was run in duplicate. The mass spectrometry proteomics data have been deposited to the ProteomeXchange Consortium via the PRIDE (57) partner repository with the dataset identifier PXD035903.

### Phospho-peptide and phospho-site identification

All raw files (.wiff) were processed to ProteinPilot software (version 4.5, SCIEX) using the Paragon algorithm (version 4. 5. 0. 0,1654). The phospho-peptides along with phospho-sites were identified against the complete sequence of Hepatitis C virus JFH-1 polyprotein. The identification settings were used as follows: (a) trypsin for proteolytic cleavage (b) Iodoacetic acid used for Cys alkylation and (c) phosphorylation emphasis used as a special factor. Peptides and proteins were validated at [1% false discovery rate (FDR) and with unused ProtScore 0.05.

## Acknowledgments

SD acknowledges the J.C. Bose Fellowship from the Department of Science and Technology (DST), Govt. of India, for research support. This study was also supported by the DBT-IISc partnership program, DST Fund for Improvement of Science and Technology Infrastructure (DST-FIST) level II infrastructure, and the University Grants Commission Centre of Advanced Studies. We also acknowledge the funding through Indo-Swiss Joint Research Program (ISJRP) from Department of Biotechnology. HR is supported by the research fellowship from the Council of Scientific and Industrial Research (CSIR-SPM). We recognize the divisional SPR facility at the Biological Science division at IISc.

---

**Figure S1.** Time-kinetics of HuR relocalisation from nucleus to cytoplasm. **(A, B)** Huh7.5 cells were infected with HCV-JFH1 virus and 12h (A) or 24h (B) post infection, cells were harvested, and nuclear cytoplasmic fractionation performed. Western blotting was done for nuclear and cytoplasmic fraction to determine the ratio of HuR in nucleus and cytoplasm. GAPDH was used as a marker for cytoplasmic fraction and Histone H3 as a marker for nuclear fraction. Densitometry was performed and the numbers in bottom indicate Nuclear to cytoplasmic ratio of HuR after normalising with H3 and GAPDH. **(C, D)** Huh7.5 cells were transfected with vector control (pCDNA3.1), WT HuR overexpression construct or S318A HuR overexpression construct as indicated. 16h post transfection, cells were infected with HCV-JFH1 virus and 48h post infection, cells were harvested, and nuclear cytoplasmic fractionation performed. Western blotting was done for nuclear and cytoplasmic fraction to determine the ratio of HuR in nucleus and cytoplasm. The lower band in HuR blot represents endogenous HuR and the upper band represents overexpressed HuR. GAPDH was used as a marker for cytoplasmic fraction and Histone H3 as a marker for nuclear fraction. Densitometry was performed for overexpressed HuR (o/e HuR) and the numbers in bottom indicate Nuclear to cytoplasmic ratio of o/e HuR after normalising with H3 and GAPDH.

**Figure S2.** Localisation of S318D HuR. **(A)** Huh7.5 cells were transfected with either Myc-WT HuR or Myc-S318D HuR and immunofluorescence staining was carried out after 48h using Alexa Fluor conjugated secondary antibodies against Myc (Red). The nucleus was counterstained with DAPI. Scale bar represents 10µm. **(B)** Nuclear and cytoplasmic ratio for Myc-HuR was quantified for images in (A) using Zen 2.3 lite software. n=10.

**Figure S3.** Huh7.5 cells were transfected with HCV-JFH1 RNA and cells harvested at indicated time-points. Western blotting was done with anti-PKCδ, and anti-actin antibodies. Appropriate HRP-conjugate secondary antibodies were used.

**Figure S4.** NS5A overexpression does not cause HuR relocalisation. Huh7.5 cells were transfected with Myc-NS5A, and immunofluorescence staining was carried out after 48h using Alexa Fluor conjugated secondary antibodies against Myc (Red) and HuR (Green). The nucleus was counterstained with DAPI. Scale bar represents 10µm.

## References

1. Lindenbach BD, Rice CM. Unravelling hepatitis C virus replication from genome to function. Nature. 2005;436(7053):933-8.

2. Jirasko V, Montserret R, Lee JY, Gouttenoire J, Moradpour D, Penin F, et al. Structural and Functional Studies of Nonstructural Protein 2 of the Hepatitis C Virus Reveal Its Key Role as Organizer of Virion Assembly. PLOS Pathogens. 2010;6(12):e1001233.

3. Ray U, Das S. Interplay between NS3 protease and human La protein regulates translation-replication switch of Hepatitis C virus. Scientific Reports. 2011;1(1):1.

4. Dentzer TG, Lorenz IC, Evans MJ, Rice CM. Determinants of the hepatitis C virus nonstructural protein 2 protease domain required for production of infectious virus. Journal of Virology. 2009;83(24):12702–13.

5. Gu M, Rice CM. Structures of hepatitis C virus nonstructural proteins required for replicase assembly and function. Current opinion in virology. 2013;3(2):129–36.

6. Niepmann M, Gerresheim GK. Hepatitis C Virus Translation Regulation. International Journal of Molecular Sciences. 2020;21(7).

7. Sharma G, Raheja H, Das S. Hepatitis C virus: Enslavement of host factors. IUBMB Life. 2018;70(1):41–9.

8. Bonamassa B, Ciccarese F, Antonio VD, Contarini A, Palù G, Alvisi G. Hepatitis C virus and host cell nuclear transport machinery: a clandestine affair. 2015;6.

9. Shwetha S, Kumar A, Mullick R, Vasudevan D, Mukherjee N, Das S, et al. HuR Displaces Polypyrimidine Tract Binding Protein To Facilitate La Binding to the 3&#x2032; Untranslated Region and Enhances Hepatitis C Virus Replication. 2015;89(22):11356-71.

10. Srikantan S, Gorospe M. HuR function in disease. Front Biosci (Landmark Ed). 2012;17(1):189–205.

11. Schultz CW, Preet R, Dhir T, Dixon DA, Brody JR. Understanding and targeting the disease-related RNA binding protein human antigen R (HuR). 2020;11(3):e1581.

12. Ma W-J, Cheng S, Campbell C, Wright A, Furneaux H. Cloning and Characterization of HuR, a Ubiquitously Expressed Elav-like Protein (*). Journal of Biological Chemistry. 1996;271(14):8144–51.

13. Marceau CD, Puschnik AS, Majzoub K, Ooi YS, Brewer SM, Fuchs G, et al. Genetic dissection of Flaviviridae host factors through genome-scale CRISPR screens. Nature. 2016;535(7610):159-63.

14. Ripin N, Boudet J, Duszczyk MM, Hinniger A, Faller M, Krepl M, et al. Molecular basis for AU-rich element recognition and dimerization by the HuR C-terminal RRM. 2019;116(8):2935–44.

15. Fan XC, Steitz JA. HNS, a nuclear-cytoplasmic shuttling sequence in HuR. 1998;95(26):15293-8.

16. Fialcowitz-White EJ, Brewer BY, Ballin JD, Willis CD, Toth EA, Wilson GM. Specific Protein Domains Mediate Cooperative Assembly of HuR Oligomers on AU-rich mRNA-destabilizing Sequences*. Journal of Biological Chemistry. 2007;282(29):20948–59.

17. Grammatikakis I, Abdelmohsen K, Gorospe M. Posttranslational control of HuR function. Wiley Interdiscip Rev RNA. 2017;8(1):10.1002/wrna.372.

18. Kim HH, Abdelmohsen K, Lal A, Pullmann R, Jr., Yang X, Galban S, et al. Nuclear HuR accumulation through phosphorylation by Cdk1. Genes Dev. 2008;22(13):1804–15.

19. Filippova N, Yang X, King P, Nabors LB. Phosphoregulation of the RNA-binding protein Hu antigen R (HuR) by Cdk5 affects centrosome function. The Journal of biological chemistry. 2012;287(38):32277–87.

20. Abdelmohsen K, Pullmann R, Jr., Lal A, Kim HH, Galban S, Yang X, et al. Phosphorylation of HuR by Chk2 regulates SIRT1 expression. Molecular cell. 2007;25(4):543–57.

21. Kim HH, Yang X, Kuwano Y, Gorospe M. Modification at HuR(S242) alters HuR localization and proliferative influence. Cell cycle (Georgetown, Tex). 2008;7(21):3371–7.

22. Lafarga V, Cuadrado A, Lopez de Silanes I, Bengoechea R, Fernandez-Capetillo O, Nebreda AR. p38 Mitogen-activated protein kinase- and HuR-dependent stabilization of p21(Cip1) mRNA mediates the G(1)/S checkpoint. Molecular and cellular biology. 2009;29(16):4341–51.

23. Liao WL, Wang WC, Chang WC, Tseng JT. The RNA-binding protein HuR stabilizes cytosolic phospholipase A2α mRNA under interleukin-1β treatment in non-small cell lung cancer A549 Cells. The Journal of biological chemistry. 2011;286(41):35499–508.

24. Doller A, Huwiler A, Müller R, Radeke HH, Pfeilschifter J, Eberhardt W. Protein kinase C alpha-dependent phosphorylation of the mRNA-stabilizing factor HuR: implications for posttranscriptional regulation of cyclooxygenase-2. Molecular biology of the cell. 2007;18(6):2137–48.

25. Doller A, Akool el S, Huwiler A, Müller R, Radeke HH, Pfeilschifter J, et al. Posttranslational modification of the AU-rich element binding protein HuR by protein kinase Cdelta elicits angiotensin II-induced stabilization and nuclear export of cyclooxygenase 2 mRNA. Molecular and cellular biology. 2008;28(8):2608–25.

26. Doller A, Schlepckow K, Schwalbe H, Pfeilschifter J, Eberhardt W. Tandem phosphorylation of serines 221 and 318 by protein kinase Cdelta coordinates mRNA binding and nucleocytoplasmic shuttling of HuR. Molecular and cellular biology. 2010;30(6):1397–410.

27. Doller A, Winkler C, Azrilian I, Schulz S, Hartmann S, Pfeilschifter J, et al. High-constitutive HuR phosphorylation at Ser 318 by PKC{delta} propagates tumor relevant functions in colon carcinoma cells. Carcinogenesis. 2011;32(5):676–85.

28. Yoon JH, Abdelmohsen K, Srikantan S, Guo R, Yang X, Martindale JL, et al. Tyrosine phosphorylation of HuR by JAK3 triggers dissociation and degradation of HuR target mRNAs. Nucleic acids research. 2014;42(2):1196–208.

29. Chu PC, Chuang HC, Kulp SK, Chen CS. The mRNA-stabilizing factor HuR protein is targeted by β-TrCP protein for degradation in response to glycolysis inhibition. The Journal of biological chemistry. 2012;287(52):43639–50.

30. Liwak-Muir U, Dobson CC, Naing T, Wylie Q, Chehade L, Baird SD, et al. ERK8 is a novel HuR kinase that regulates tumour suppressor PDCD4 through a miR-21 dependent mechanism. Oncotarget. 2016;7(2):1439–50.

31. Filippova N, Yang X, Nabors LB. Growth factor dependent regulation of centrosome function and genomic instability by HuR. Biomolecules. 2015;5(1):263–81.

32. Vigouroux C, Casse JM, Battaglia-Hsu SF, Brochin L, Luc A, Paris C, et al. Methyl(R217)HuR and MCM6 are inversely correlated and are prognostic markers in non small cell lung carcinoma. Lung cancer (Amsterdam, Netherlands). 2015;89(2):189–96.

33. Zhou HL, Geng C, Luo G, Lou H. The p97-UBXD8 complex destabilizes mRNA by promoting release of ubiquitinated HuR from mRNP. Genes Dev. 2013;27(9):1046–58.

34. de Sousa GF, Lima Mde A, Custodio DF, Freitas VM, Monteiro G. Chemogenomic Study of Carboplatin in Saccharomyces cerevisiae: Inhibition of the NEDDylation Process Overcomes Cellular Resistance Mediated by HuR and Cullin Proteins. PLoS One. 2015;10(12):e0145377.

35. Wang W, Fan J, Yang X, Fürer-Galban S, Lopez de Silanes I, von Kobbe C, et al. AMP-activated kinase regulates cytoplasmic HuR. Molecular and cellular biology. 2002;22(10):3425–36.

36. Wang W, Yang X, Kawai T, de Silanes IL, Mazan-Mamczarz K, Chen P, et al. AMP-activated Protein Kinase-regulated Phosphorylation and Acetylation of Importin α1: INVOLVEMENT IN THE NUCLEAR IMPORT OF RNA-BINDING PROTEIN HuR*. Journal of Biological Chemistry. 2004;279(46):48376–88.

37. Kumar R, Mehta D, Mishra N, Nayak D, Sunil S. Role of Host-Mediated Post-Translational Modifications (PTMs) in RNA Virus Pathogenesis. Int J Mol Sci. 2020;22(1).

38. Neveu G, Barouch-Bentov R, Ziv-Av A, Gerber D, Jacob Y, Einav S. Identification and Targeting of an Interaction between a Tyrosine Motif within Hepatitis C Virus Core Protein and AP2M1 Essential for Viral Assembly. PLOS Pathogens. 2012;8(8):e1002845.

39. Dehghani B, Hashempour T, Hasanshahi Z, Moayedi J. Bioinformatics Analysis of Domain 1 of HCV-Core Protein: Iran. International Journal of Peptide Research and Therapeutics. 2020;26(1):303–20.

40. Kato K, Yamanouchi D, Esbona K, Kamiya K, Zhang F, Kent KC, et al. Caspase-mediated protein kinase C-delta cleavage is necessary for apoptosis of vascular smooth muscle cells. Am J Physiol Heart Circ Physiol. 2009;297(6):H2253–H61.

41. Xie J, Lee JA, Kress TL, Mowry KL, Black DL. Protein kinase A phosphorylation modulates transport of the polypyrimidine tract-binding protein. Proc Natl Acad Sci U S A. 2003;100(15):8776–81.

42. Otsuka M, Kato N, Moriyama M, Taniguchi H, Wang Y, Dharel N, et al. Interaction between the HCV NS3 protein and the host TBK1 protein leads to inhibition of cellular antiviral responses. Hepatology (Baltimore, Md). 2005;41(5):1004–12.

43. Lu L, Zhang Q, Wu K, Chen X, Zheng Y, Zhu C, et al. Hepatitis C virus NS3 protein enhances cancer cell invasion by activating matrix metalloproteinase-9 and cyclooxygenase-2 through ERK/p38/NF-κB signal cascade. Cancer letters. 2015;356(2 Pt B):470-8.

44. Borowski P, Oehlmann K, Heiland M, Laufs RJJov. Nonstructural protein 3 of hepatitis C virus blocks the distribution of the free catalytic subunit of cyclic AMP-dependent protein kinase. 1997;71(4):2838-43.

45. Borowski P, zur Wiesch JS, Resch K, Feucht H, Laufs R, Schmitz HJJoBC. Protein kinase C recognizes the protein kinase A-binding motif of nonstructural protein 3 of hepatitis C virus. 1999;274(43):30722–8.

46. Meng Z, Liu Q, Sun F, Qiao L. Hepatitis C virus nonstructural protein 5A perturbs lipid metabolism by modulating AMPK/SREBP-1c signaling. Lipids in Health and Disease. 2019;18(1):191.

47. Mankouri J, Tedbury PR, Gretton S, Hughes ME, Griffin SDC, Dallas ML, et al. Enhanced hepatitis C virus genome replication and lipid accumulation mediated by inhibition of AMP-activated protein kinase. Proc Natl Acad Sci U S A. 2010;107(25):11549–54.

48. Wu C-H, Hsu F-T, Chao T-L, Lee Y-H, Kuo Y-C. Revealing the suppressive role of protein kinase C delta and p38 mitogen-activated protein kinase (MAPK)/NF-κB axis associates with lenvatinib-inhibited progression in hepatocellular carcinoma in vitro and in vivo. Biomedicine & Pharmacotherapy. 2022;145:112437.

49. Wang J, Guo Y, Chu H, Guan Y, Bi J, Wang B. Multiple Functions of the RNA-Binding Protein HuR in Cancer Progression, Treatment Responses and Prognosis. 2013;14(5):10015–41.

50. Papatheofani V, Levidou G, Sarantis P, Koustas E, Karamouzis MV, Pergaris A, et al. HuR Protein in Hepatocellular Carcinoma: Implications in Development, Prognosis and Treatment. Biomedicines. 2021;9(2):119.

51. Chen T, Dai X, Dai J, Ding C, Zhang Z, Lin Z, et al. AFP promotes HCC progression by suppressing the HuR-mediated Fas/FADD apoptotic pathway. Cell Death & Disease. 2020;11(10):822.

52. Wu M, Tong CWS, Yan W, To KKW, Cho WCS. The RNA Binding Protein HuR: A Promising Drug Target for Anticancer Therapy. Current cancer drug targets. 2019;19(5):382–99.

53. Jiang X, Tan H-Y, Teng S, Chan Y-T, Wang D, Wang N. The Role of AMP-Activated Protein Kinase as a Potential Target of Treatment of Hepatocellular Carcinoma. Cancers (Basel). 2019;11(5):647.

54. Yang X, Liu Y, Li M, Wu H, Wang Y, You Y, et al. Predictive and preventive significance of AMPK activation on hepatocarcinogenesis in patients with liver cirrhosis. Cell death & disease. 2018;9(3):264-.

55. Sadria M, Seo D, Layton AT. The mixed blessing of AMPK signaling in Cancer treatments. BMC Cancer. 2022;22(1):105.

56. Holmes B, Benavides-Serrato A, Freeman RS, Landon KA, Bashir T, Nishimura RN, et al. mTORC2/AKT/HSF1/HuR constitute a feed-forward loop regulating Rictor expression and tumor growth in glioblastoma. Oncogene. 2018;37(6):732–43.

57. Perez-Riverol Y, Bai J, Bandla C, García-Seisdedos D, Hewapathirana S, Kamatchinathan S, et al. The PRIDE database resources in 2022: a hub for mass spectrometry-based proteomics evidences. Nucleic acids research. 2022;50(D1):D543–D52.

