## Supplementary figures and images for "Hepatitis C virus non-structural proteins modulate cellular kinases for increased cytoplasmic abundance of host factor HuR and facilitate viral replication"

### Supplementary Fig 1

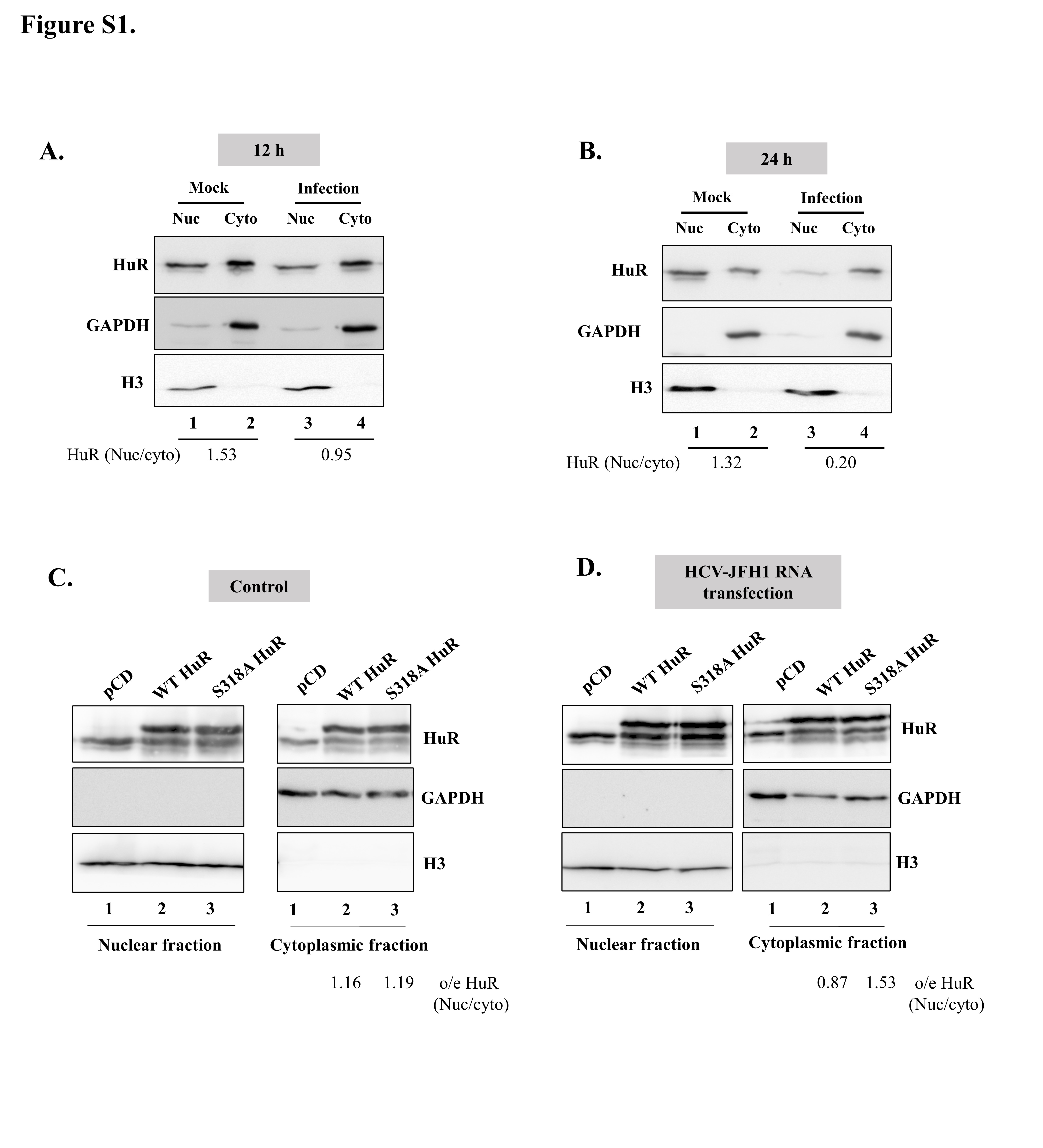

### Supplementary Fig 2

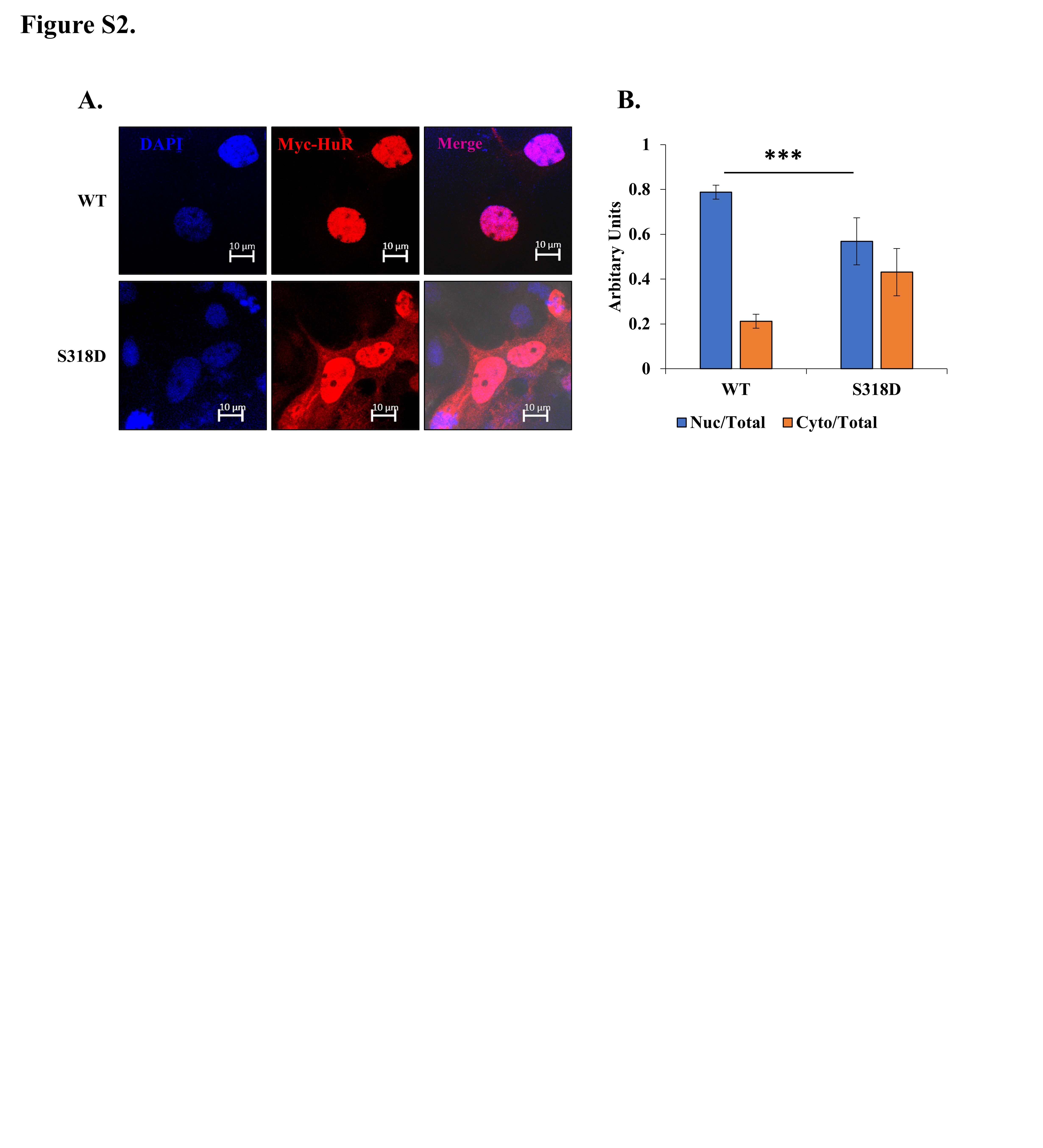

### Supplementary Fig 3

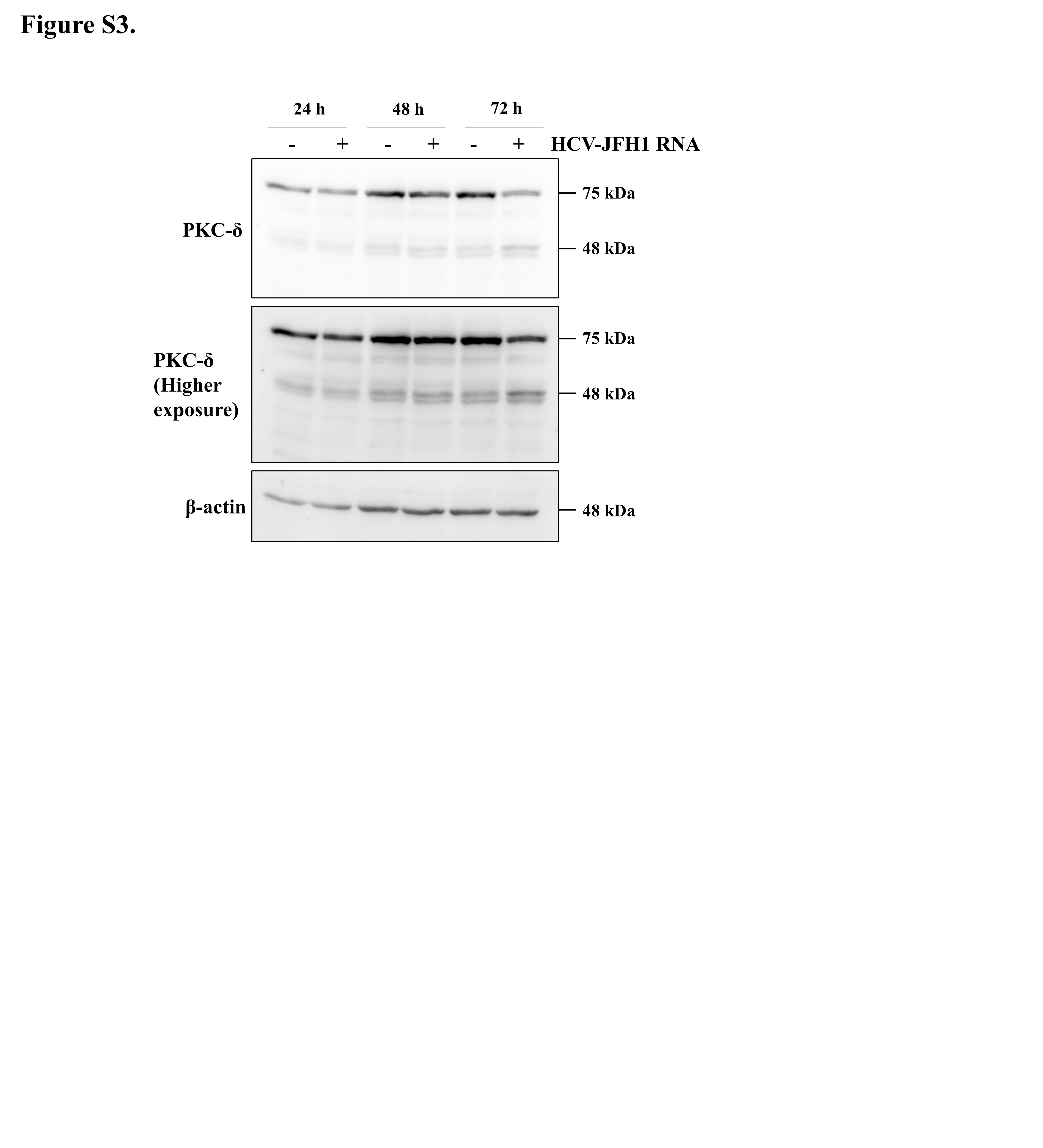

### Supplementary Fig 4

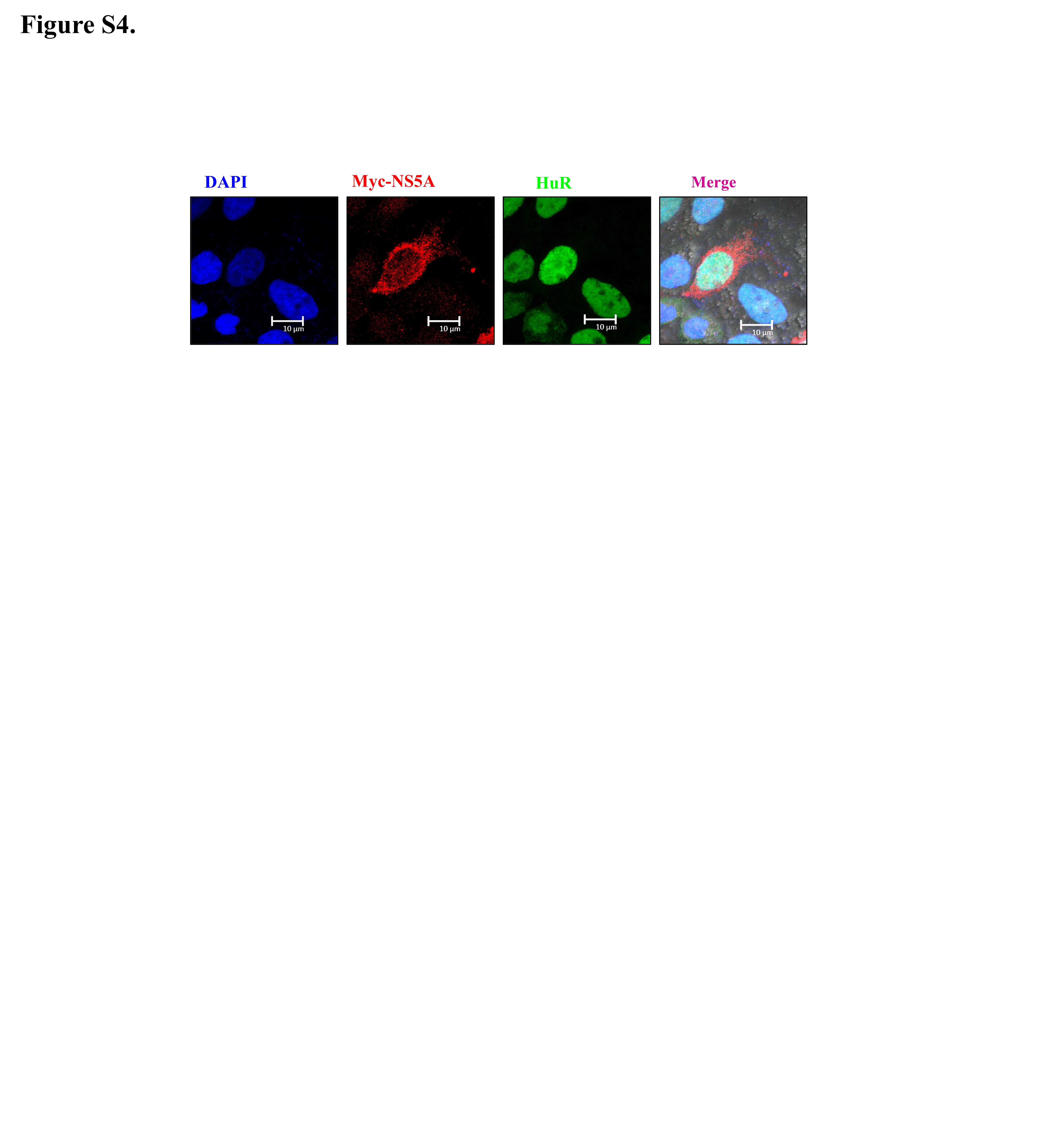
